# Resurrecting Golgi proteins to grasp Golgi ribbon formation and self-association under stress

**DOI:** 10.1101/2021.07.27.453980

**Authors:** Luis F. S. Mendes, Mariana R. B. Batista, Emanuel Kava, Lucas Bleicher, Mariana C. Micheletto, Antonio J. Costa-Filho

**Author notes:** To whom correspondence should be addressed: Antonio J. Costa-Filho, DF/FFCLRP/USP, Av. Bandeirantes 3900, Ribeirão Preto, SP, Brazil, 14040-901.

## Abstract

The Golgi complex is a membranous organelle located in the heart of the eukaryotic secretory pathway. A subfamily of the Golgi matrix proteins, called GRASPs, are key players in the stress-induced unconventional secretion, the Golgi dynamics during mitosis/apoptosis, and Golgi ribbon formation. The Golgi ribbon is vertebrate-specific and correlates with the appearance of two GRASP paralogs (GRASP55/GRASP65) and two coiled-coil Golgins (GM130/Golgin45), which interact with each other *in vivo*. Although essential for the Golgi ribbon formation and the increase in Golgi structural complexity, the molecular details leading to their appearance only in this subphylum are still unknown. Moreover, despite the new functionalities supported by the GRASP paralogy, little is known about the structural and evolutionary differences between these paralogues. In this context, we used ancestor sequence reconstruction and several biophysical/biochemical approaches to assess the evolution of the GRASP structure, flexibility, and how they started anchoring their Golgin partners. Our data showed that the Golgins appeared in evolution and were anchored by the single GRASP ancestor before *gorasp* gene duplication and divergence in Metazoans. After the *gorasp* divergence, variations inside the GRASP binding pocket determined which paralogue would recruit each Golgin partner (GRASP55 with Golgin45 and GRASP65 with GM130). These interactions are responsible for the protein’s specific Golgi locations and the appearance of the Golgi ribbon. We also suggest that the capacity of GRASPs to form supramolecular structures is a long-standing feature, which likely affects GRASP’s participation as a trigger of the stress-induced secretory pathway.

## Introduction

The classical secretory pathway is the most important delivery system in eukaryotic cells. It is responsible for carrying a nascent protein from inside the endoplasmic reticulum (ER) towards the Golgi complex and then its final destination. This orchestrated transport was first described by the Nobel Prize winner George Palade [^1^], and since then, it has been a subject of intense studies. Surprisingly, although this *ER-Golgi-Final Destination* pathway is the best known and characterized secretion pathway in eukaryotes, it is not the only one [^2^]. Recently, many proteins were observed to be transported through pathways that do not require some, if any, of the classical machinery components [^2^]. There is still not a clear common feature to all these alternative delivery systems, but they have been all together classified as Unconventional Protein Secretion (UPS) pathways.

A possible common point for these UPS pathways seems to be its stress-triggered mechanism, including nutrient starvation [^3^], temperature [^4^], and ER stress [^5^]. Some proteins targeted to UPS are translocated into the ER but somehow travel to their final destinations bypassing the Golgi complex (Type IV UPS) [^6^,^7^]. Other proteins are soluble and dispersed in the crowded cytosol but recruited to the UPS pathway to perform additional functionalities outside the cell without the direct participation of the ER or the Golgi (Type III UPS) [^3^,^8^]. Others are directly transported across the membrane by specific ABC transporters (Type II UPS) or by pore-mediated translocation (Type I UPS). Moreover, there might still be other pathways for protein delivery, illustrating the magnificent complexity of protein transport in eukaryotic cells [^2^]. UPS in eukaryotes is an emergent theme, and several proteins that are directly transported by these pathways have been implicated in distinct excretion mechanisms and multiple human illnesses, including cystic fibrosis [^6^], Alzheimer’s disease [^9^], and diseases arising from inflammation [^10^].

A common point between Types III and IV UPS and the classical secretory pathway is the involvement of a Golgi matrix protein family called Golgi Reassembly and Stacking Proteins (GRASPs) [^11-14^]. GRASPs constitute a family of peripheral membrane-associated proteins, which were first suggested as an essential factor in the Golgi cisternae reassembly after mitotic times [^15,16^]. GRASPs are observed in all the branches of the eukaryotic tree of life, although not equally spread in all supergroups [^17^]. In Metazoans, evolution led to the appearance of two GRASP paralogue genes called *gorasp1* and *gorasp2* (translating the proteins GRASP65 and GRASP55, respectively) [^18^]. GRASP65 is anchored to the cell membrane through the myristoylation of its glycine-2. The interaction with a protein partner, GM130, seems to determine its orientation at the membrane surface [^19,20^]. This direct interaction was suggested to be the main responsible for the preferential *cis*-Golgi attachment of GRASP65 [^19^]. On the other hand, GRASP55 can be myristoylated and palmitoylated *in vivo* and is localized in the *medial*/*trans*-Golgi through the interaction with Golgin45 [^21^]. GRASP55 and 65 are necessary to anchor GM130 and Golgin45 to the Golgi. Moreover, these four proteins are also essential for the formation and integrity of the Golgi ribbon [^22,23^]. Although membrane anchoring has proved essential for GRASP tethering, the general occurrence of myristoylation in eukaryotes besides mammalians remains obscure.

GRASPs are structurally organized in two main portions. The N-terminal one, called the GRASP domain (DGRASP), is formed by two PDZ subdomains with an unusual circular permutation in eukaryotes, making them resemble prokaryotic PDZs [^24^]. The PDZs are connected in tandem to allow rigid body reorganization, facilitating their interaction with multiple different protein partners [^25^]. This promiscuity for interaction with several different protein partners and their unusual structural plasticity [^26,27^] might be the convergent features of GRASPs in different cell functional tasks, including as anchoring factors of Golgins [^12,15,23^], their direct participation in Golgi dynamics during mitotic and apoptotic times [^28-30^] and their self-association during stress conditions triggering UPS [^4,31^]. It comes as quite a surprise that there is still a significant gap in understanding how GRASPs can perform those functionalities. The second portion of the GRASP’s structure, the so-called Serine and Proline-Rich (SPR) domain, has a regulatory function [^15^] and presents a very low sequence identity even for closely related species [^25^]. Despite the lack of experimental data on the structure of the SPR, previous computational analyses suggested it as fully disordered for most systems [^26^].

One current debate about GRASPs concerns whether they are essential for Golgi cisternae stacking [^23^]. Rothman and co-workers observed that several different tethering factors synergically work in the Golgi organization [^32^], thus suggesting that GRASPs share the protagonism in the Golgi maintenance with other proteins. On the other hand, upon cell treatment with brefeldin A, the Golgi membrane and its oligosaccharide-modifying enzymes relocate to the ER. In contrast, the Golgins and GRASPs accumulate in the cytoplasm keeping their ability to form a ribbon-like reticulum [^33^]. This suggests that Golgins and GRASPs would be sufficient for the Golgi structuration. Furthermore, Golgins have recently been shown to undergo liquid-liquid phase separation when overexpressed [^34,35^] and GRASPs to form amyloid-like fibrils [^36^]. Those pieces of information suggest that the participation of the Golgi-matrix proteins in the Golgi organization might be revisited. More specifically, how would the synergy between GRASPs and the other Golgi matrix proteins facilitate the Golgi organization, and, considering that the Golgi is an ancient organelle, how old would this type of self-association be?

The complexity of life lies in the fate of evolution. The first eukaryotic species to arise in history is still a matter of debate. However, estimates for the Last Eukaryote Common Ancestor (LECA) age range from 1–1.9 billion years ago [^37^]. The tree of life spreads through different kingdoms, and each branch differentiates itself so that the phenotype similarities are somehow hard to identify in most cases. Nevertheless, the genotype still carries some traces of that first eukaryote representative. Interestingly, although a Golgi with differentiated compartments and trafficking pathways directly relates to the Eukaryogenesis, GRASPs are one of the few Golgi-matrix proteins predicted to be present in LECA [^38^]. Here, we used ancestor sequence reconstruction and several biophysical/biochemical tools to gain insight into GRASP ancient proteins and compare them with modern human relatives. The GRASP ancestors were used to understand three points that are fundamental for Golgi ribbon formation and cell-stress responses in UPS: when and how GRASPs started recruiting their Golgins to the Golgi; how the GRASP structure has evolved to make them a hub in the protein interactome; and how disorder, flexibility, and self-association (under stress) have developed in the modern human orthologues.

## Results

### Resurrecting ancient proteins to unravel GRASP history

Our phylogenetic analyses started by collecting GRASP orthologues in several branches of the eukaryotic tree and predicting the corresponding GRASP domain for each one of them. We excluded the disordered SPR region from the analyses because of its low sequence identity, even in closely related species [^12,26^]. For the ancestor sequence reconstruction, a high-quality sequence alignment is mandatory. After the analyses, the GRASP tree with the highest probability reflected the different separations of clades of eukaryotes (Figure S1), thus showing the robustness of the constructed evolutionary tree. The first reconstructed node of the GRASP history was the connection between the GRASP55 and GRASP65 inside the Metazoa kingdom (Figure 1A). This first ancestor (ANC1) would represent the GRASP from the first Metazoan to appear in the evolution, based on the used methodology and database. The second ancestor (ANC2) was built based on the node connecting the Metazoa with the Fungi kingdom, suggesting a GRASP from the first representative of the Opisthokonta group (Figure 1A). The third ancestor (ANC3) was built based on the connection of the subgroup used to reconstruct the ANC2 with representatives from the Amoebozoa taxonomic group (Figure 1A). ANC3 would be a GRASP sequence representing the approximate point of the Unikonts/Bikonts divergence. The ancestor four (ANC4) was built based on the whole group used to construct ANC3 plus representatives from the SAR+Haptophyta (Figure 1A). This was the closest to LECA reconstructed with reasonable statistics for the predicted protein sequence (Figure S1). The sequence alignment between the ancestors and the modern human DGRASP55/65 suggested that all the ancestors kept the characteristic two-PDZ fold of DGRASPs (Figure 1B). In fact, the AlphaFold 2 (DeepMind) software retrieves models for the GRASP ancestors where the structural superposition with the crystallographic structures of GRASP55 and GRASP65 have a Cα RMSD of less than 2.5 Å (Figure S2).

**Figure 1.**
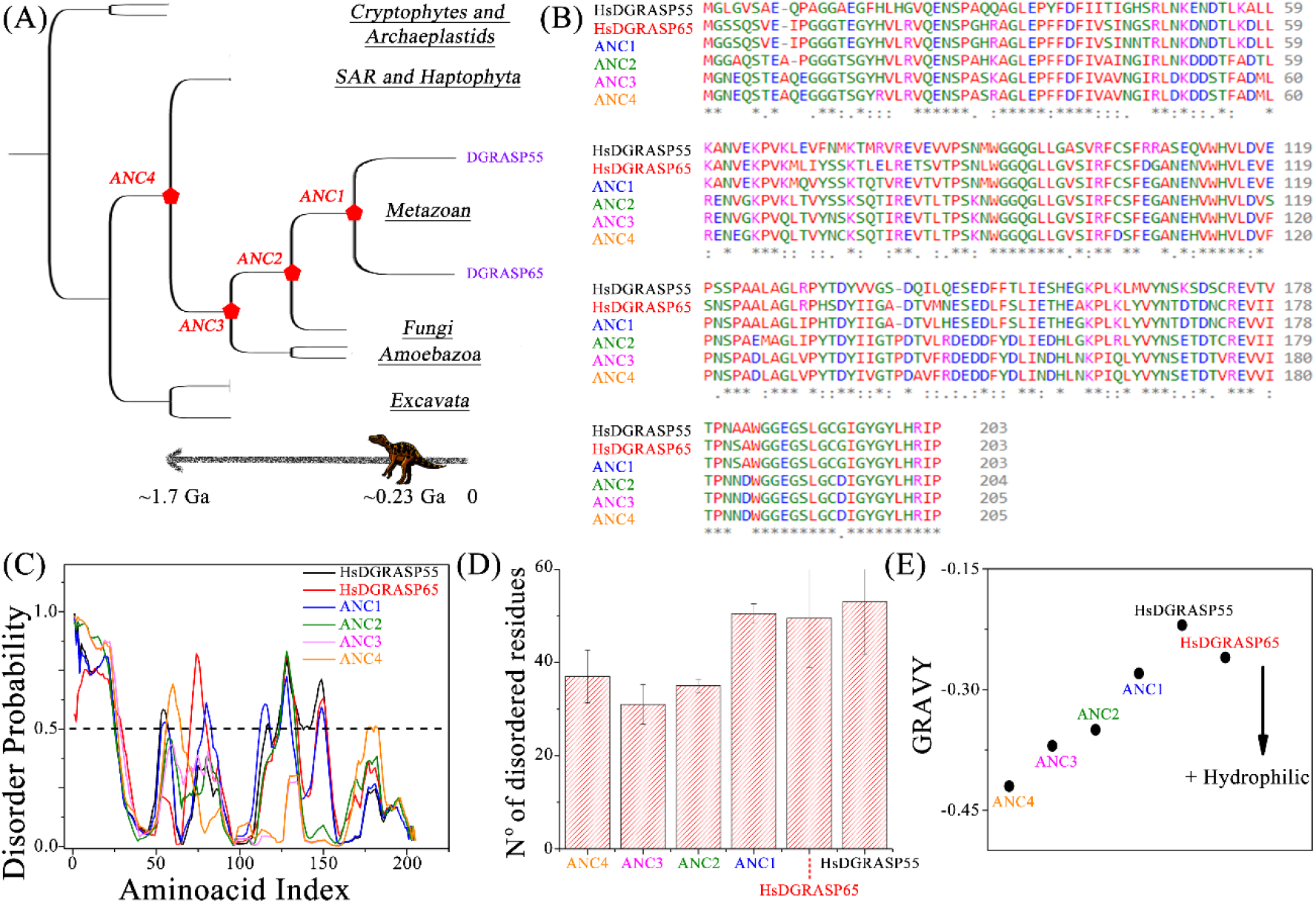
Phylogenetic and hydrodynamic characterization of the ancestor proteins compared with the modern orthologues. (A) Phylogenetic tree of the eukaryotes highlighting the points where the theoretical ancestor sequences would have appeared in evolution. (B) Sequence alignment between the reconstructed ancestors and the modern human GRASP55 and GRASP65 sequences. (C) Disorder predictions using PONDR VXLT. A sequence with a final score higher than 0.5 is considered disordered. (D) The mean number of residues predicted to be in intrinsically disordered regions calculated for each modern and ancestor protein. The standard error is considered as a combination of the outputs of the PONDR VXLT, CAN_XT, and VS2L software. (E) The grand average of hydropathy (GRAVY) value for each modern and ancestor protein was determined as the sum of hydropathy values of all the amino acids divided by the number of residues in the sequence [^39^]. The hydrophobicity of a protein increases with the GRAVY value.

The functional promiscuity of GRASPs has already been correlated with their high structural flexibility in solution, primarily due to the massive presence of intrinsically disordered regions (IDR) in their structures [^17,26^]. However, it has been experimentally demonstrated that, in Metazoans, the presence of IDRs is a particular hallmark of DGRASP65 [^17^]. This raised the hypothesis that the ancestor of GRASP55/65 should have characteristics similar to GRASP65, especially regarding the presence of IDRs and the structural promiscuity at the tertiary structure level [^17^]. We then repeated the search for IDRs in the ancestors to understand the presence of IDRs throughout the evolution of GRASPs. Contrary to what we expected, our data indicated a somewhat lower number of residues predicted to be in IDRs in ancient proteins, suggesting that disorder is a modern feature (Figures 1C and D). However, these differences were rather related to the extension of the conserved IDRs than to the appearance of new IDR regions during evolution (Figure 1C).

Going deeper into the disorder prediction, we analysed the ancestors and modern GRASP dynamics via predictions for the fast backbone movements directly from their amino acid sequences using DynaMine [^40^]. For all the tested proteins, a significant portion of the DGRASP core was classified as having “highly context-dependent dynamics”, suggesting that the cores can change depending on the physicochemical context where those proteins are inserted (Figure S3) [^41^]. Therefore, a significant portion of the DGRASP core is malleable, and its overall dynamic is adjusted depending on the environment, a property that goes back to the beginning of GRASP history.

Finally, the overall content of amino acids also suggested a tendency to decrease the total hydrophobicity. The grand average of hydropathy (GRAVY), calculated by adding the hydropathy value for each residue and dividing it by the length of the sequence [^42^], increased from the last ancestor to the human paralogues (Figure 1E). This indicates a decrease in protein solubility with evolution.

### The distribution of coevolving residues and frustration explains Metazoa’s GRASP-Golgin specificity

The GRASP55/65 protein family presented a pattern of residue coevolution in which many highly conserved residues coevolved (to be referred to as Set 1, with 66 residues), together with seven smaller coevolving sets presenting 8, 7, 4, 3, 3, 2 and 2 residues, which are limited to specific sequences in this protein family. Set 1 included the residues involved in zinc binding (Cys102/103 and His17/18 in rat GRASP65/GRASP55), present in ANC1-3 but not in ANC4. In the latter, there was an arginine instead of the histidine and an aspartate instead of the cysteine, suggesting the zinc-binding site was not yet available in the early evolution of the GRASP55/65. ANC4 could present a salt bridge instead, as suggested in its structural model (Figure S4).

The larger coevolving set also included most residues involved in the binding to the Golgi matrix protein GM130 (His20, Trp113, Met164, Leu152, Gly138, His18, Val137, Leu143, Cys103, Arg101, Asp140, Val100, Ile142, Ser99, Ala98, Leu96, Gly97, and Gly94) and those involved in dimerization (Leu116, His200, Leu199, Tyr198, Arg201, Tyr196, Gly195, Gly188, Ile194, and Cys192). About half of the residues involved in GM130 recognition were present in all four ancestral sequences. This fact may explain why GRASP55 can bind GM130 *in vitro*, although with a much lower affinity [^21^]. A similar picture was found when analysing the prevalence of residues for the interaction with Golgin45, whose binding interface shared many of the residues seen for the complex with GM130 (numbering refers to *Mus musculus* GRASP55, PDB ID 5H3J). Residues His18, Leu20, Ile37, Gly94, Leu96, Gly97, Val98, Ser99, Ile100, Arg101, Phe102, Cys103, Tyr164 and Arg174, necessary for the binding between GRASP55 and Golgin45, were all present in the larger set. Since GM130 and Golgin45 are Holozoa-specific proteins (present only in Metazoans and their single-celled relatives) and precede GRASP55/65 duplication (which is Metazoa-specific) [^38^], their first appearance fits the expected time for ANC1. The conservation of residues found in Set 1 would thus entail ANC1 to bind to GM130 and Golgin45. This was indeed found when we measured ANC1 binding to the mammalian GM130 and Golgin45 C-terminal region. The affinity strengths were similar to the values observed in the interaction of human GRASP55/65 with their respective Golgins (Figure 2A).

**Figure 2:**
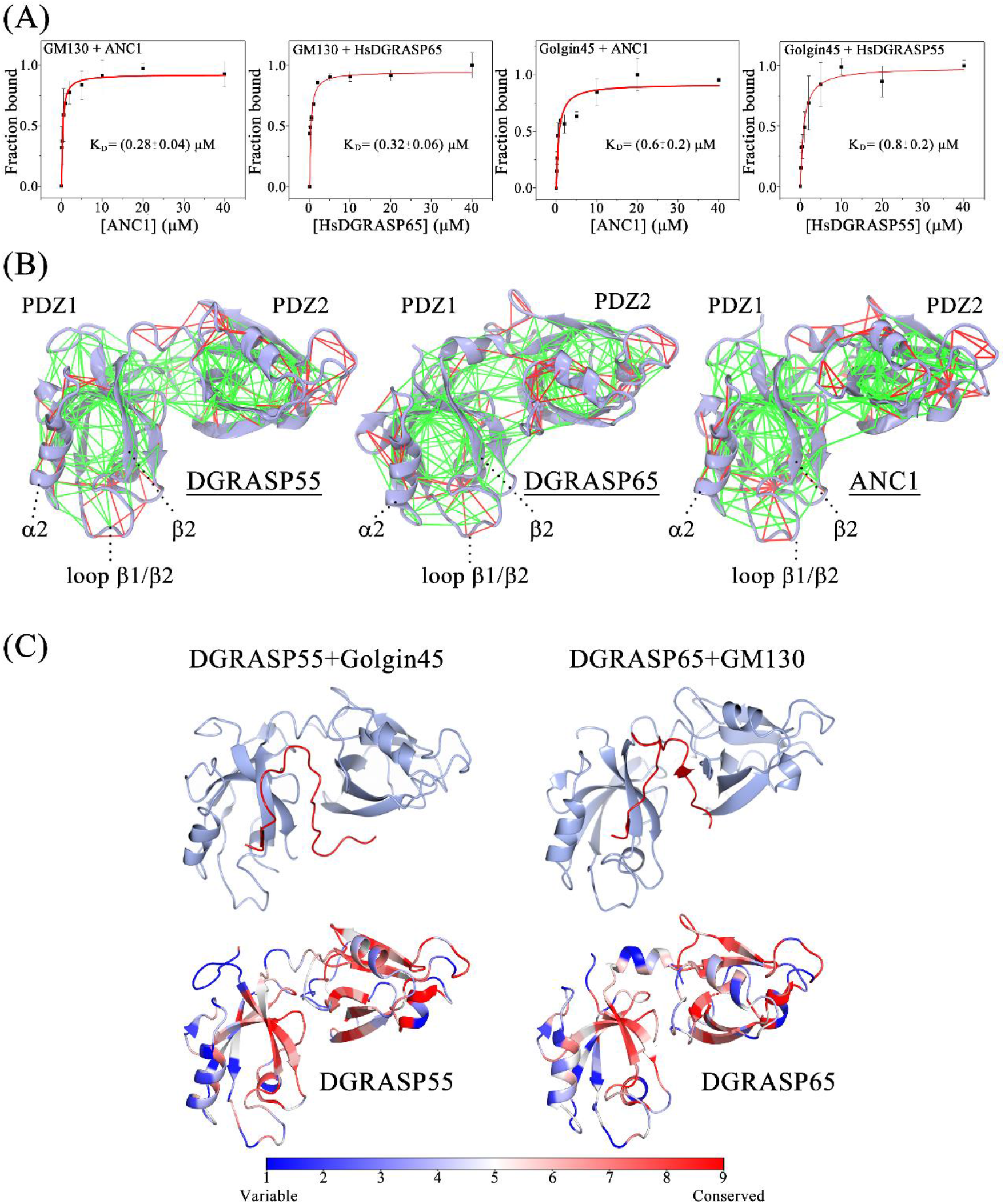
GRASP-Golgin specificity along with evolution and the history of the PDZ1 binding pocket. (A) Binding isotherms of ANC1, DGRASP55, DGRASP65 with the Abz-labelled human GM130 and Golgin45 C-terminus region. A “One binding site” model was fitted to the data using OriginPro 8.5 (OriginLab Corporation). Error bars represent standard deviations of triplicate measurements. (B) Configurational frustration patterns of DGRASP55, DGRASP65, and ANC1 were calculated using Frustratometer [^47^]. The colour scale indicates the local mutational frustration where the minimally (green) and highly frustrated (red) sites are shown. (C) Ribbon representation (in light blue) of mammalian DGRASP55 and DGRASP65 bound to the C-terminal region of Golgin45 and GM130 (red), respectively (PDB IDs 5H3J and 4REY). The structures were constructed using CCP4MG. The structure-mapped sequence conservation of both GRASPs is also presented. Surface representations of both GRASP55 and GRASP65 were coloured according to the degree of evolutionary conservation based on a dataset of 300 homologs collected from Uniprot for each protein (identity between homologs varying from 30 to 95%) using ConSurf [^48^]. Blue indicates high sequence variability, and red indicates high sequence conservation.

The evolution of the GRASP65/GM130 and GRASP55/Golgin45 complex formation had a milestone: the appearance of GM130 and Golgin45. Right after that appearance in the divergence of Holozans, the existing GRASP ancestor (ANC1) recruited the Golgins to the Golgi. The data in Figure 2A and the set of coevolving residues that came back from the LECA show that the molecular determinants of such recruitment were already present in GRASPs much before the appearance of those Golgins. In a second moment, GRASP paralogues appeared and evolved to the situation where GRASP55 (GRASP65) would recruit only Golgin45 (GM130). The overall conservation of the GRASP binding region in both Golgins shows that these regions did not diverge along with evolution (Figure S5). Therefore, it was the GRASPs that differentiated with time to recruit just one of the Golgins each. The high structural similarity of the GRASP’s PDZs [^17^] indicates that the amino acid content and the relative orientation of the PDZs determined the GRASP-Golgin specificity.

Another interesting conservation is found after the analyses of the protein’s frustration patterns. Sites with low local configurational frustration (green lines in Figure 2B) usually populate the hydrophobic core of folded proteins [^43^]. Maximally frustrated linkages (red lines in Figure 2B) often indicate biologically relevant regions, such as those involved in binding. For instance, frustrated local networks co-localize with regions implicated in forming oligomeric interfaces [47] and regions involved in allosteric transitions [^44^]. GRASPs and their ancestors showed a conserved high degree of frustration in the interactions within *α*_2_ helix and the loop connecting *β*_1_ and *β*_2_ (Figure 2B and S6). The strand *β*_2_ and the helix *α*_2_ form the binding groove of GRASP’s PDZs [^12,45,46^]. Interestingly, the GRASP promiscuous interactome was previously suggested as a direct outcome of the somewhat low stability and high flexibility of the *α*_2_ helix in the PDZ1, a region enriched in high configurational frustration (Figure 2B) [^25^].

The recruitment of GM130 and Golgin45 by GRASP65 and GRASP55, respectively, involves the PDZ1 binding pocket and the surface area of the cleft between PDZ1 and 2 (Figure 2C). The degree of frustration observed in the cleft between PDZ1 and PDZ2 is not similar when GRASP55 and GRASP65 structures are compared, showing that GRASP65 behaves more like the ancestor ANC1 (Figure 2B). When the degree of evolutionary conservation was mapped onto the GRASP55 and GRASP65 structures, the binding pocket of PDZ1 showed the highest variability, particularly in the *α*_2_ helix and its surroundings (Figure 2C). This coincided with the maximally frustrated region seen in that helix (Figure 2B). Therefore, even though ANC1 could recruit both Golgins to the Golgi apparatus, evolution determined which GRASP paralogue would recruit which Golgin based on a differential evolutionary pressure acting in the PDZ1 binding pocket and the surface cleft between PDZ1 and PDZ2.

### Evolution made GRASPs less thermal/chemical/pH resistant

We further explored the ancestor structures by monitoring their overall stability in some physiologically relevant conditions. Firstly, the thermodynamic profiles of protein unfolding were studied using differential scanning calorimetry (DSC). The ancestors showed greater thermal stability and a tendency of increasing such stability over time regression (Figure 3A). DGRASP65 had the lowest T_M_ value and the broadest unfolding transition, reflecting its lowest structural stability and the low cooperativity of unfolding. The low cooperativity of the thermal transition comes from the lower number of tertiary contacts in the DGRASP65 structure, a property already discussed elsewhere [^17^]. DGRASP55 was more thermally stable than DGRASP65 with a thermal transition at 47°C, which was still considerably lower than ANC1 (T_M_∼62°C), ANC2 (T_M_∼73°C), ANC3 (T_M_∼74°C) and ANC4 (T_M_∼77°C). Interestingly, we found a linear dependence when the apparent T_M_ for each ancestor was plotted as a function of the estimated time when the ancestor organism lived (Figure 3B).

**Figure 3:**
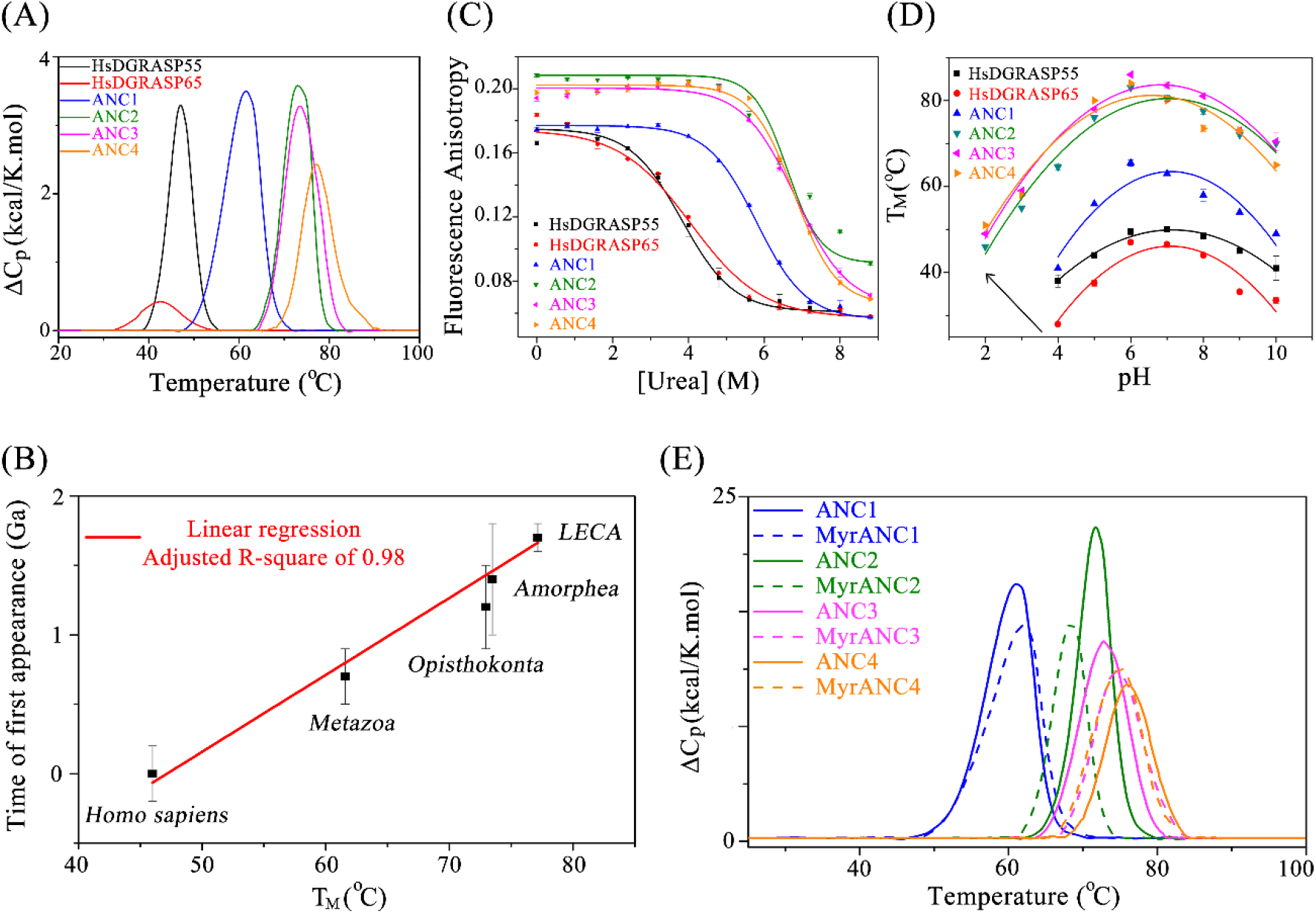
Thermodynamic analyses of the ancestor proteins compared with the modern human orthologues. (A) Excess heat capacity of the modern and ancestor proteins at a fixed protein concentration. (B) The experimental T_M_ values obtained in the DSC curves are plotted against the theoretical time of the first appearance estimated from the evolution point where the ancestors were expected to be present, including the variances. The references for the estimated time of first appearances are given in: Homo sapiens [^49^], Metazoa [^50^], Opisthokonta [^37^], Amorphea [^37^], and LECA [^51^]. Error bars were taken as the uncertainty in the estimated times of the first appearance given in the references and should not be considered as well-established values. (C) Chemical denaturation using the chaotropic agent urea as monitored by fluorescence anisotropy. Lines are fits of the Boltzmann equation to the experimental data. (D) Dependence of the T_M_ values as a function of the solution pH monitored by the fluorescence changes of the extrinsic probe SYPRO Orange using Differential Scanning Fluorimetry. Lines are fits of a second-degree polynomial function to the experimental data. (E) Excess heat capacity of the ancestor proteins and their myristoylated version at a fixed protein concentration and in the presence of detergent (0.03% DDM). Error bars represent standard deviations of duplicate (D) and triplicate (C) measurements.

Chemical perturbations, monitored by fluorescence anisotropy, and pH variations, monitored by Differential Scanning Fluorimetry (DSF), showed the same increase in structural stability for the ancestors (Figures 3C and D). In the chemical perturbation, since the tryptophan residues are conserved (Figure 1B), the fluorescence anisotropy values in the native condition are good indicators of structural flexibility in those regions. Hence, the higher values of anisotropy observed in the absence of urea indicate an increase in the ancestral proteins’ structural rigidity compared to the human orthologs.

### Mammalian GRASPs are myristoylated in vivo, and so are their ancestors

Prediction of N-terminal myristoylation, a well-established post-translational modification in mammalian GRASP, by neural networks using Myristoylator [^52^] showed that the four GRASP ancestors have a medium-to-high propensity of being myristoylated *in vivo* (Table S1). To confirm that the ancestors could indeed be myristoylated, we used a strategy based on the *in vitro* myristoylation of the protein’s N-terminus performed by an N-Myristoyl transferase [^53,54^]. The first indirect observation of the success of the myristoylation strategy was the significant decrease in the protein solubility, likely due to the hydrophobic tail located in its N-terminus. Purification of myristoylated proteins required the presence of 0.03% DDM. The same buffer solution was used to purify the non-myristoylated versions for comparison (see Materials and Methods).

Our circular dichroism (CD) data indicated that the myristoylated tail did not disturb the secondary structure content (Figure S7). However, it did affect the protein stability for all the cases tested (Figure 3E). Nevertheless, the effect did not follow a regular pattern of alterations in parameters such as T_M_ or the calorimetric enthalpy variation, with increases and decreases observed that were not dependent on the time in evolution (Table S2). The only pattern followed by nearly all the proteins after myristoylation seemed to be the decrease in the cooperativity of the unfolding transition as estimated by the ΔT_1/2_ (Table S2). Our data strongly support the hypothesis that, akin to the human paralogues, all the ANC proteins could be myristoylated *in vivo*. Such protein modification is likely an old trend in the GRASP family.

### Amyloid-like aggregation tendency and GRASPs as stress sensors

The increase in structural flexibility and the decrease in protein stability observed in the modern orthologues could allow GRASPs to undergo conformational changes with an impact on cell function. For instance, human GRASPs and the GRASP orthologue in *S. cerevisiae* have been observed to form amyloid-like aggregates [^55,4,36^]. Other reports have shown that several Golgin proteins undergo liquid-liquid phase separation (LLPS), including GM130, golgin160, GMAP210, and 31, golgin97, golgin245, GCC88, and GCC185 [^34,35^]. Based on the apparent tendency of higher-order structure formation in some of the Opisthokonta family of Golgi-matrix proteins and the close relation between LLPS and protein fibrillation [^56-60^], we next addressed how old this fibrillation tendency would be in the GRASP family.

In that context, we expected that, should higher-order structure formation be necessary for a particular GRASP functionality during normal or cell-stress conditions, it must have always happened along with the evolution. Since fibrillation and any other type of self-association are a concentration-dependent phenomenon, we decided to study the ancestor’s thermal denaturation as a function of the protein concentration. The unfolding transition curves should be concentration-independent unless oligomerization or aggregation processes take place. The DSC traces for the ANCs at low concentration (1 mg/mL) were monomodal and typical of a two-state transition (Figure 4). The DSC curves shifted toward higher temperatures (increase in apparent T_M_) and became bimodal (Figure 4) upon concentration increase. In this bimodal DSC trace, the peak centred at the higher T_M_ seemed to become dominant as protein concentration raised. In the case of ANC1, the transition at the highest concentration presented a single endothermic peak (Figure 4). A similar tendency was also observed for the ANC 2, 3, and 4 with their DSC traces becoming gradually monomodal but without reaching a single peak stage. A single transition would likely be present in these cases should a higher concentration be used, which was not possible due to solubility issues.

**Figure 4.**
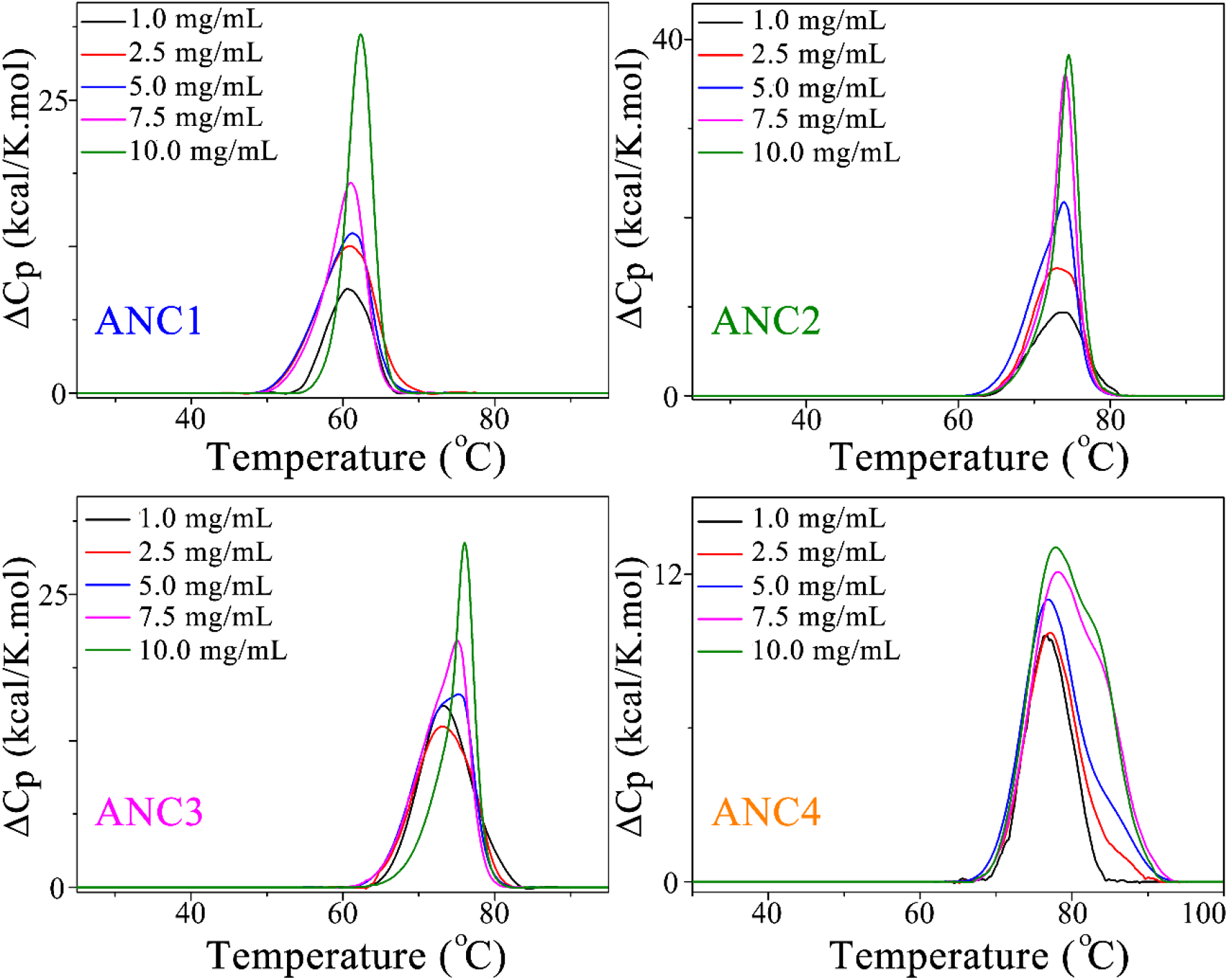
Excess heat capacity of the ancestor proteins using different protein concentrations (1.0, 2.5, 5.0, 7.5, and 10.0 mg/mL) in buffer A. The raw DSC traces were subtracted with the buffer baseline and then normalized by protein concentration.

Our data indicated the existence of a concentration-dependent self-association process, which increased the ancestors’ thermal stability. Given that aggregation is a concentration-dependent process and that some GRASP representatives can form amyloid fibres, the samples from the DSC run at the highest concentration were tested for the typical autofluorescence of amyloid fibres (Figure 5A) [^61^]. Just like observed for Aβ40, K18 tau, γ-B-crystallin, HEWL, and I59T lysozyme [^62,63^], all the ancestor proteins showed the classical intrinsic fluorescence in the visible range that is a signature of amyloid formation (Figure 5A-B). The ancestor’s fibres also showed the classical increase in affinity for the amyloid-detector dye ThT (Figure 5C). Collectively, our data support the conclusion that, just like the modern orthologues in *Homo sapiens* and yeast, GRASPs have an amyloid-like formation propensity that comes from LECA and that is concentration- and temperature-dependent.

**Figure 5:**
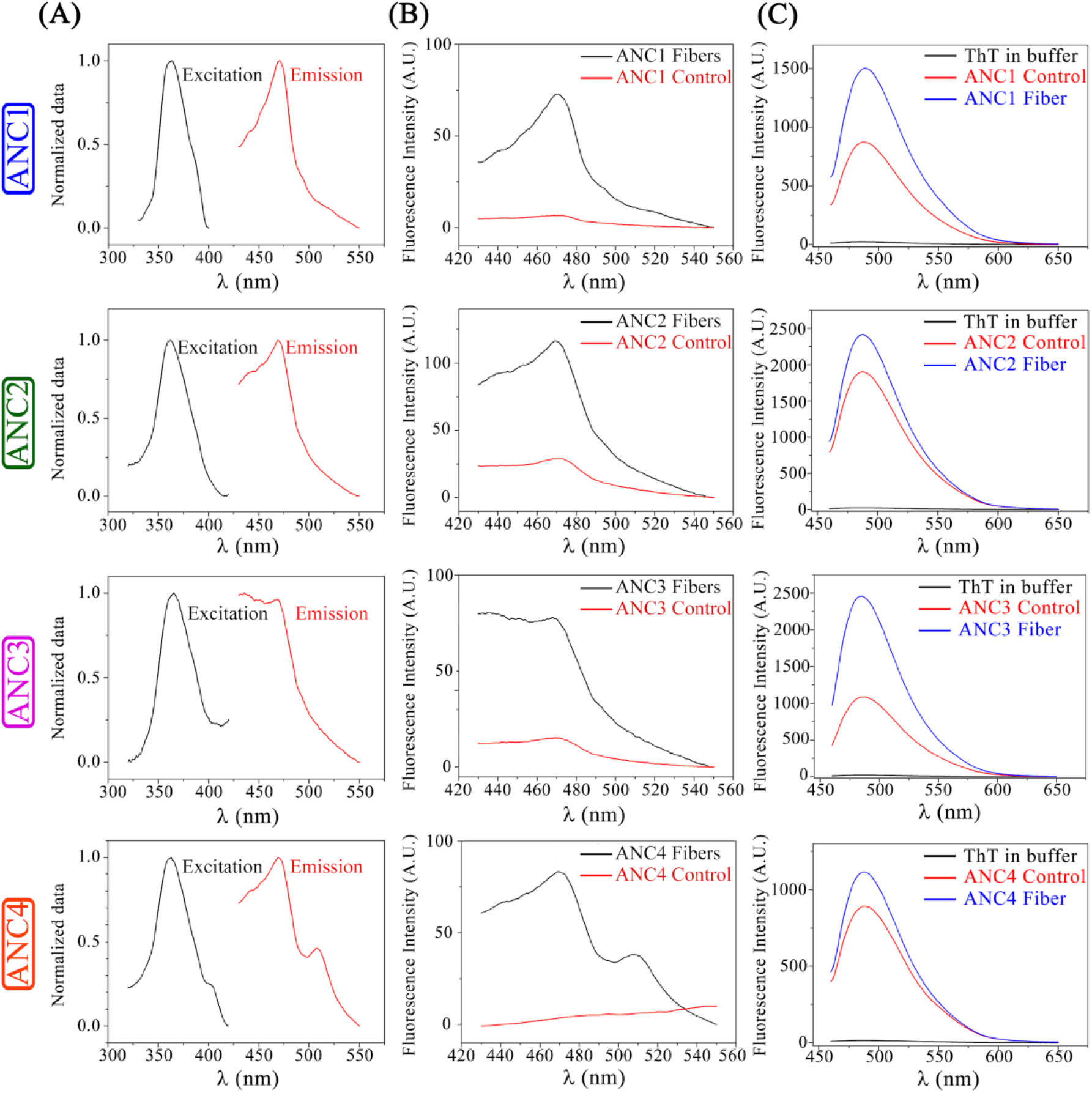
Spectroscopic properties of the thermally induced aggregation observed for each ancestor protein after the DSC experiments. The ancestors were tested for the excitation/emission peaks characteristic of the amyloid fibres (column 1). The emission spectra using the excitation fixed at 365 nm were collected for the proteins before and after the thermal treatment (column 2). The thermally treated samples were also tested for the increase in affinity for ThT (column 3) compared with the protein solution before the thermal treatment and for the ThT fluorescence in the absence of protein in the buffer solution.

To show that only a mild denaturation condition was necessary for the fibrillation process to happen, the kinetics of the fibrillation process was monitored measuring the intensity of the autofluorescence from the amyloid fibres (Figure 6). Using a temperature value high enough to increase the rate of amyloid formation but still significantly below the ancestor T_M_ values (Figure 4A), we showed that all the ancestors had a nucleation-dependent fibrillation curve that was completely absent in the control sample kept at room temperature (Figure 6). The ANC1 was the ancestor with the slowest kinetics, taking a lag time of more than one day before reaching full saturation. The ANC2, on the other hand, was the fastest, reaching a saturation profile in a couple of minutes (Figure 6). It is not clear the reasons for this significant difference in the fibrillation kinetics. However, the time interval for the ANC2 fibrillation agrees with what was previously observed for the *S. cerevisiae* GRASP [^4^]. Since the ANC2 is predicted to be the ancestor of the Metazoans with the Fungi kingdom, it seems that the faster fibrillation tendency was maintained in the Fungi examples but strongly decreased in the Metazoans. In agreement with this observation, it has been shown that the saturation of the curves for the human GRASPs needed a couple of days to occur [^36^]. ANC3 and ANC4 had comparable kinetics, although ANC4 fibrillated faster and in a more cooperative manner. Our data suggest that GRASPs have always been fibrillation-prone proteins, and the increase of this aggregation tendency is somehow controlled depending on the branch of the eukaryotic tree.

**Figure 6:**
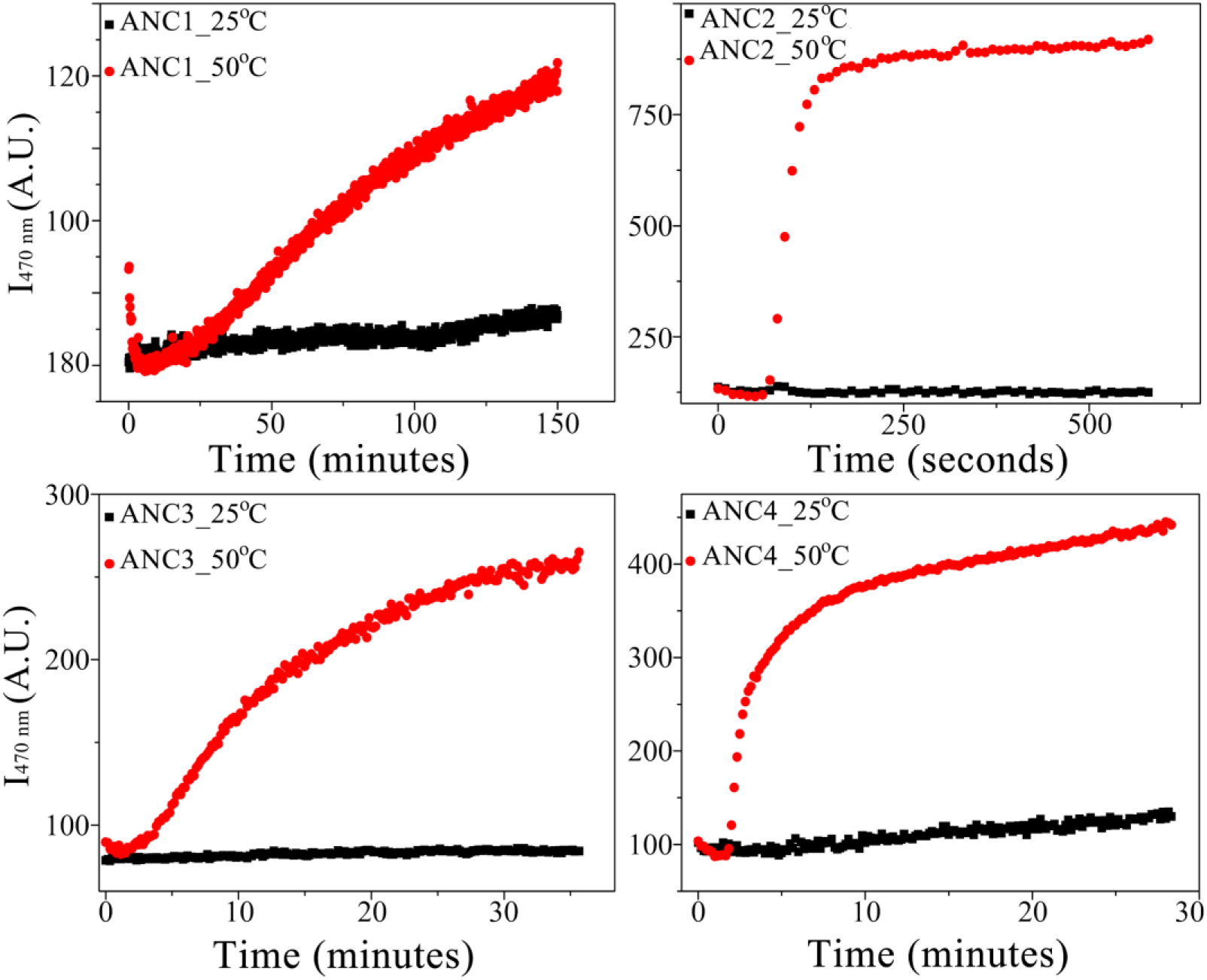
Fibrillation kinetics of the ancestor proteins monitored by the fibre autofluorescence intensity at 470 nm as a function of time. It is worth noting that 50°C is considerably lower than the ancestor’s T_M_ values (Figure 4A).

To further explore other conditions involved in triggering self-association under thermal stress besides concentration, we fixed the protein concentration and varied the ionic strength of the solution. Figure S8 shows the DSC traces, and once more, the curves became bimodal as the salt concentration was raised. To show that the higher salt concentration was not inducing self-association without the structural perturbation caused by the temperature, room-temperature SEC-MALS experiments at high ionic strength were performed (Figure S9). The data clearly showed that all the ANCs, akin to the human paralogues, kept the monomeric organization at room temperature. The higher ionic strength had no effect without the increase in temperature (Figure S9). Therefore, the ancestor’s aggregation process, just like it was observed before for the human and fungi GRASPs [^4,36,55^], is indeed stress-induced (thermal stress in the situation tested here) and controlled by the protein concentration and the ionic strength of the solution.

To shed light on why an external perturbation is necessary for self-association, an aggregation analysis was performed using the CamSol method [^64,65^]. The method relies on two different analyses: one based solely on the primary protein sequence and a structure-based method of calculating a solubility profile, which accounts for the proximity of the amino acids in the 3D structure and their solvent exposure [^65^]. It is important to note several hotspots for protein aggregation when only the amino acid sequence is analyzed (intrinsic analyses), seen in all the ancestor proteins (Figure 7). However, several of those hotspots of aggregation vanished when Camsol considered the protein tertiary structure (Figure 7). Therefore, a perturbation of the protein structure is pivotal to exposing the sites for aggregation and inducing the fibrillation process, which was achieved in the DSC/fluorescence experiments by increasing the temperature (Figures 5 and 6). In this sense, it was previously observed that the GRASP orthologue in *S. cerevisiae* formed amyloid fibres only at a temperature close to its T_M_ or in acidic solutions [^4^]. The activation of GRASP fibrillation could be related to a protective mechanism in non-permissive temperatures. The same profile was also observed *in vivo* during nutrient starvation, suggesting remarkable plasticity of function for these aggregates [^31^]. It is worth pointing that the human GRASPs contain more aggregation hotspots in their GRASP domain compared with the SPR [^36,55^]. The GRASP in *S. cerevisiae* was also shown to fibrillate even in the absence of the SPR region [^4^]. Therefore, it seems that the GRASP domain drives GRASP fibrillation in modern systems as well as in their ancestor’s counterparts.

**Figure 7:**
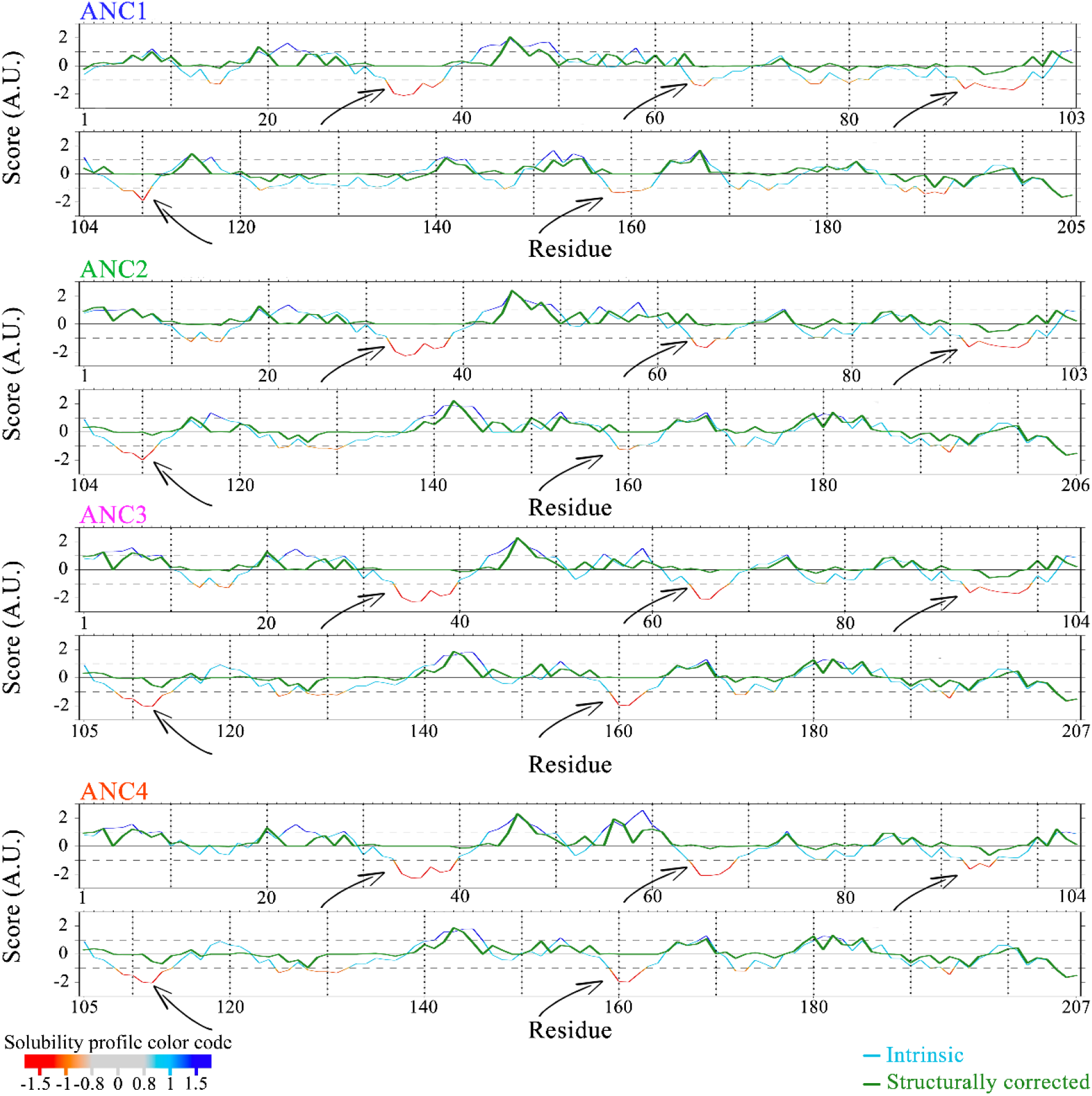
CamSol analyses of the ancestor sequences based on the intrinsic prediction from the primary sequence and the structurally corrected prediction using the predicted structure from molecular modelling. The (structurally corrected) solubility score from –1.5 to 1.5 is mapped on the protein’s amino acid sequences and represented in a colour gradient from red (low solubility) to blue (high solubility). The arrows indicate the regions with the lowest solubility scores.

## Discussion

GRASPs form a family of malleable proteins with unique structural properties [^11,12,14,17,25,26^], several different functionalities [^11^,^15^] and a promiscuous interactome [^25^]. Despite being central hubs in the cell interactome, the detailed description of their roles in those different processes still deserves more attention. The number of genes codifying GRASP among organisms can vary. Plants do not have any obvious GRASP/GM130/Golgin45 homologues but have stacked Golgi [^18^]. Fungi, on the other hand, have one GRASP homolog, which anchors a GM130-like protein called Bug1 (not a GM130 homolog), and their Golgi varies from stacked to fully dispersed in the cytosol [^26^]. Mammals have two GRASPs located in different positions in their stacked Golgi. A better understanding of how GRASP structural properties have evolved can certainly impact the description of the diverse set of cell processes involving that Golgi matrix protein. Among those processes, the Golgi structure organization, the out-of-normal conditions observed in cell response to stress (particularly those that trigger UPS), and the many Golgi perturbations observed in diseases and apoptosis are of special interest [^30,66,67^].

A first attempt to correlate the biophysical properties of GRASPs from different organisms showed that GRASP65 was more like GRASPs from lower eukaryotes [^17^]. Here, we used modern computational and molecular biology methods to turn on the time machine in protein evolution, therefore making it possible to follow aspects of the protein’s biochemical and biophysical properties over time. We showed that going backward in time, GRASPs were more rigid, thermally and chemically stable, and tolerant to pH variations (Figure 8A). Evolution seems to have pressed GRASP to be more flexible by increasing the extension of its IDRs and decreasing its overall tertiary structural stability, flexibility, and compaction (Figure 9A). Therefore, it is curious to note that, upon the appearance of the Metazoa paralogy GRASP55/GRASP65 in evolution, the tendency of higher flexibility was kept mainly by GRASP65. This indicates that GRASP55 is suffering from a unique evolutionary pressure and is probably involved in Metazoa-specific functionalities. It has been observed that GRASP55 is more involved in UPS [^13,68^], energy sensing in the Golgi [^69^], and membrane tethering in autophagy under cell stress [^70^]. GRASP65 participates in Golgi dynamics, especially in processes such as Golgi organization [^71,72^], Golgi fragmentation in apoptosis [^73^], and modulation of Golgi structure and microtubule organization during cell division [^74,75^]. Therefore, it is likely that GRASP65’s higher flexibility and similarity with lower eukaryotes [^17^] are determinants of its participation in Golgi dynamics. On the other hand, participation in UPS seems to require a more “ordered” and less malleable structure as GRASP55.

**Figure 8:**
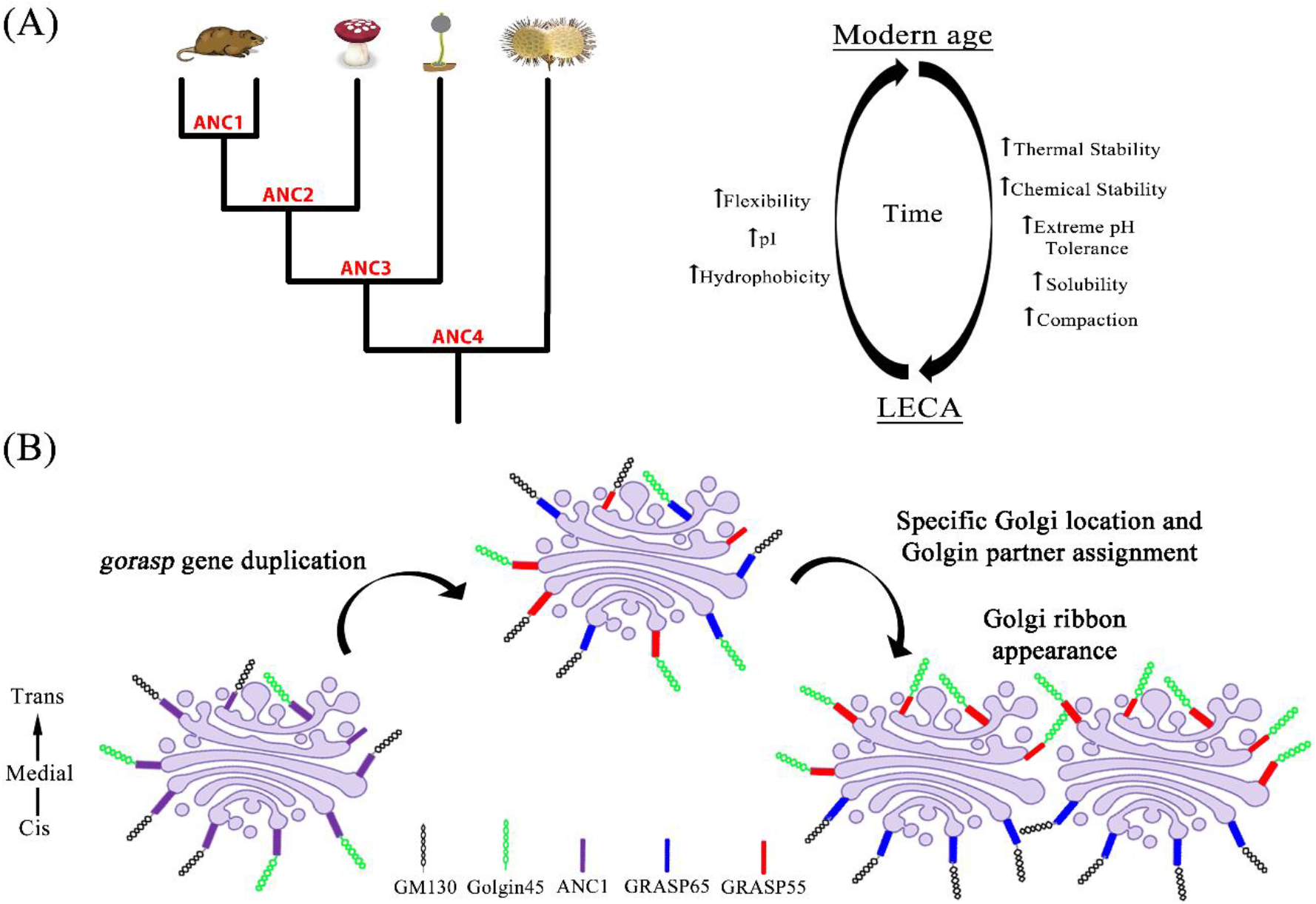
Lessons about GRASP coming from their ancestors. (A) Biophysical properties that GRASPs acquired during evolution and those significantly altered. (B) A model for the origin of the Golgi ribbon in vertebrates. At first, before the appearance of Golgin45 and GM130 in holozoans, the ancient Golgi was equally populated by a GRASP homolog here represented as the ANC1. Right after the appearance of Golgin45 and GM130, our data suggest that the ancestor GRASP could anchor both Golgins with high affinity. After the *gorasp* gene duplication, both proteins started to diverge so that they differentiate in Golgi location and golgin partners: GRASP65 with GM130 in the *cis*-Golgi, and GRASP55 binds Golgin45 in the *trans*/*medial* faces. Since the coordinated action of these four proteins is necessary for the Golgi ribbon formation, that would be the birth of the Golgi ribbon.

Regarding their domain constituents, the GRASP domain (DGRASP) has been under the spotlight much more frequently than the disordered and highly variable SPR domain. Within DGRASP, the once believed as fully ordered PDZ sub-domains have also shown different biophysical properties [^17^]. The further dissection of the PDZ’s biophysical properties showed that, despite the PDZ1 variability, the glycine in position 2 was fully conserved in the ancestor’s sequences (Figure 1B). We demonstrated that, in the presence of a broad-spectrum N-myristoyltransferase and the myristate substrate, all the ANCs were successfully N-myristoylated in our *in vivo* model, suggesting that, given the proper conditions, they could also suffer such modification in their native extinct organisms. Myristoylation is an old protein modification that plays several different roles in eukaryotic systems. The binding to the membrane directly relates to GRASP functionalities, including its oligomerization tendency and Golgi organization [^11^].

Therefore, we expect that the GRASP ancestors were already associated with the Golgi complex membranes in the past (Figure 8B).

Interestingly, we showed here that ANC1, the common ancestor of GRASP55 and GRASP65, interacted with both GM130 and Golgin45 with high affinity (Figure 2A). It is worth noting that our estimated time for ANC1 existence roughly coincides with the appearance of Golgin45 and GM130. Therefore, when Golgin45 and GM130 appeared in evolution, an ancestor GRASP (ANC1) capable of recruiting both Golgins was already bound to the Golgi complex, probably without any preferential Golgi location (Figure 8B). When the *gorasp* gene duplication (GRASP65-Chromosome 3 and GRASP55-Chromosome 2) occurred, there was no reason for any preferential binding between these new GRASPs and the Golgins (Figure 8B). Hence, the specific assignment of the GRASP binding partners and their location in the Golgi came after an evolutionary pressure, yielding the pair GRASP65/GM130 in the *cis*-Golgi, and the GRASP55/Golgin45 in the *trans/medial* faces. In both cases, the pairs of proteins were particularly close to the rims of the Golgi stacks [^23^] (Figure 8B). This is directly reflected in the degree of sequence variability observed inside the binding pocket of PDZ1 and in the overall degree of frustration on the cleft between PDZ1 and PDZ2 (Figure 2). Since the coordinated action of these four proteins is necessary for the Golgi ribbon formation [^22,23^], this might have been the birth of the Golgi ribbon (Figure 8B).

Another intriguing feature of modern GRASPs is their capacity to form higher-order structures under stress [^4^,^36^]. Other Golgi-related proteins, including GM130, have shown involvement in forming exquisite compartments via LLPS [^34,35^]. The intimate relationship between those two processes led us to explore the fibrillation phenotype along with evolution. Our data indicate that GRASP has always been a stress sensor capable of adjusting itself to mediate protein-protein interactions (PPIs) and form complex structures, like amyloid-like fibrils. It is impacting to notice how most of those features were preserved along with evolution and how they could have naturally driven GRASPs to perform several of their different cell functionalities.

We then propose that GRASPs could act in two different cell scenarios. In scenario 1, when native conditions were present, GRASPs would work as tethering factors capable of mediating several different PPIs essential for the correct flow of the exocytic pathway. They would also participate in the formation of the higher-order architecture of the Golgi in synergy with the Golgins [^14,22,23^]. The latter property might have been the natural cause of GRASP duplication since GRASP65 is responsible for retaining GM130 and p115 into the *cis*-Golgi. At the same time, GRASP55 in the *trans*/medial Golgi faces is responsible for the retention of Golgin45 [^22^]. The need for two GRASPs in Metazoans is also justified by the unique participation of these proteins in the Golgi ribbon formation (a vertebrate-specific structural organization [^76^]) and as a negative regulator of exocytic transport, necessary for correct protein-cargo glycosylation [^77^]. In scenario 2, under stress, GRASPs could form higher-order structures with a potential cell-protective functionality and with direct involvement in UPS. Stress conditions were shown to induce GRASP phosphorylation [^2,68^] and O-GlcNAcylation [^78^], then breaking its oligomerization properties and also its retention in the Golgi membranes. Therefore, GRASP could work as tethering factors in UPS and be involved in forming condensates or aggregated structures, recruiting specific proteins for UPS. GRASP fibrillation under stress, both chemically (starvation-triggered) and under heat, was already observed *in vivo* and shown to be reversible [^31^].

In conclusion, we demonstrated here that GRASP constitutes a family of myristoylated proteins with high structural plasticity that can respond to stress conditions by inducing a self-association process, which might be relevant in UPS. Remarkably, their promiscuous interactome correlates with significant variability in the binding pocket of PDZ1 and an extensive array of conserved highly frustrated interactions in the surface of both PDZs. We showed that all these features are not restricted to the modern mammalian GRASP but seem to be already present in GRASPs back to LECA. More than that, these proteins adapted themselves to the many variations of Earth’s temperature along with evolution without losing their structural plasticity. How this adaptation specifically impacts the plethora of functions performed by GRASPs remains to be unravelled. Several exciting themes are therefore still open for future studies involving this moonlighting family of Golgi-matrix proteins.

## Materials and Methods

### Ancestor sequence reconstruction (ASR)

To resurrect the GRASP ancestors, the sequences of the GRASP orthologues were manually collected using the primary sequence of human GRASP65 and GRASP55 as templates in the NCBI database. The GRASP domain for each orthologue was predicted using Pfam [^79,80^]. An initial database was constructed comprising >100 DGRASP orthologues, and the first refinement was performed by sequence alignment using ClustalΩ [^81^]. The quality of the alignment is essential for the Ancestor Sequence Reconstruction (ASR). All the sequences with unique insertions/deletions were manually removed. A total of 83 DGRASP orthologues were used in the final analyses. The sequences were aligned with MUSCLE v3.8.31 [^82^] and MSAProbs 0.9 [^83^]. Phylogeny was inferred using a maximum likelihood method implemented in RAxML [^84^]. These processes were interactively performed using Phylobot [^85^]. Phylobot searches for the tree and branch lengths with the highest probability of producing the sequence alignment based on a collection of Markov models of amino acid substitution [^85^]. Therefore, the software builds ML trees for all combinations of sequence alignments and evolutionary models in its collection. Ancestral protein sequences were reconstructed using the empirical Bayes approach as implemented in the software CODEML [^86^] controlled by Lazarus [^87^]. The final computational data can be found in http://www.phylobot.com/portal/status/VnJfz/. The ancestors with the highest probability estimated in the sequence alignment method and Markov model were synthesized by GeneScript with codon optimization for *E. coli* heterologous expression and subcloned into a pET28a (Novagene) vector using Nco1 and Xho1 restriction sites.

### Protein Expression and Purification

The human DGRASP55 and DGRASP65 were expressed and purified as described elsewhere [^17^]. The protocol for the ancestor proteins production was the same as the one used for modern human proteins. The final purified protein solutions were 20 mM Tris/HCl, 150 mM NaCl, 5 mM 2-Mercaptoethanol, pH 8.0 (Buffer A). GRASP myristoylation was performed using an adapted protocol described in [^53,54^]. Due to the low solubility of the myristoylated protein, the working buffer for these proteins was 1X PBS + 150 mM NaCl, 5 mM 2-Mercaptoethanol, 0.03% DDM, pH 7.4. Comparisons between myristoylated and non-myristoylated proteins were always made with the proteins in the same working buffer.

### Fluorescence spectroscopy

Steady-state fluorescence was monitored using a Hitachi F-7000 spectrofluorometer equipped with a 150 W xenon arc lamp and polarized filters for anisotropy experiments. The excitation and emission monochromators were set at 5 nm slit width. For tryptophan fluorescence anisotropy, the samples were selectively excited at 300 nm. The anisotropy values were calculated as the mean average of the values from ±10 nm (1 nm step acquisition) over the wavelength of maximum emission determined for each condition. All experiments were performed at 25 °C. The anisotropy experiments were performed in triplicate.

The isotherms of binding between ANC1 and the human C-terminus mimicking peptides of GM130 (Abz-GSNPCIPFFYRADENDEVKITVI) and Golgin45 (Abz-HPYTRYENITFNCCNHCRGELIAL) were performed at a fixed temperature of 25°C. Peptides were chemically synthesized by Biomatik with a purity by HPLC of >95%. The experiments were performed by fixing the peptide amount at 1 µM and varying the ANC1, DGRASP55, or DGRASP65 concentrations, with an overnight incubation time at 4°C without agitation. Fluorescence measurements were performed using an excitation of 350 nm, data collection from 370 to 600 nm, and excitation/emission monochromators at 10 nm slit width. The experiments were done in triplicate.

Amyloid fibre autofluorescence measurements were performed using an excitation of 365 nm and the spectra collected from 430-550 nm. The excitation and emission monochromators were set at 10 nm slit width. Excitation peaks were collected by monitoring the fluorescence intensity at 470 nm using the same parameters. In the Thioflavin (ThT) fluorescence, the fluorophore was kept at a concentration of 20 µM in all the conditions tested. Details of the sample preparation for these experiments are described in the DSC section. All experiments were repeated at least twice.

For the fibrillation kinetic experiments, 5 mg/mL of protein solutions were inserted in the equipment previously equilibrated at 25°C or 50°C, and the fluorescence intensity at 470 nm (excitation at 365 nm) was collected over time using a 10-second step. A cuvette containing each ancestor protein was first tested at a temperature of 25°C (control sample). The same protein solution/cuvette was then removed, and the fluorimeter was warmed up to 50°C using a water bath. The sample was added to the machine, and the kinetic experiment was started immediately after. All experiments were repeated at least twice.

### Differential scanning fluorimetry (DSF)

Protein melting temperature (T_M_), assuming a two-state transition model, was determined by monitoring the fluorescence intensity variation as a function of temperature for the extrinsic probe SYPRO Orange (Invitrogen). The assays were performed in an Agilent Mx3005P qPCR System equipped with a FAM SYBr green I filter with excitation and emission wavelengths of 492 nm and 516 nm, respectively. The thermal variations were in the range 25-95°C in a stepwise increment of 1 °C/minute step and using a 96-well PCR plate (Agilent Technologies) sealed with optical quality sealing tape (Microseal® ‘B’ seal from BIO-RAD). Data were analysed using the software Origin 8.5. The fluorescence intensity as a function of the temperature was individually analysed for the classical two state-transition shapes. The T_M_ was determined as the maximum intensity point of the first-order derivative curve. For the experimental setup, buffer solutions from pH 2 to 10 were prepared and filtered using a 0.2 µm filter paper: 50 mM of glycine/HCl (pH 2.0-3.0), 50 mM sodium acetate/acetic acid (pH 4.0), 50 mM sodium phosphate (pH 5.0– 8.0), and 50 mM glycine NaOH (pH 9.0 and 10.0). Protein solutions were concentrated to 2 mg/mL, and a dilution of 10 times was prepared in each buffer. All the DSF experiments were performed in duplicate.

### Differential Scanning Calorimetry (DSC)

The experiments were performed in a Nano-DSC II—Calorimetry Sciences Corporation, CSC (Lindon, Utah, USA). A heating rate of 1°C/minute was used to sweep from 25-100°C at a controlled pressure of 3 atm. The reversibility of all transitions was tested by collecting repeated heating scans. To explore the thermotropic effects caused by varying the protein concentrations, measurements were carried out in 20 mM Tris/HCl, 150 mM NaCl, 5 mM 2-Mercaptoethanol, pH 8.0 with protein concentration ranging from 1-10 mg/mL. To explore the effects of the ionic strength on the thermotropic behaviour of the ancestors, a fixed protein concentration of 2.5 mg/mL was used in a 20 mM Tris/HCl, 5 mM 2-Mercaptoethanol, pH 8.0 with the NaCl concentrations of 0 (< 1 mM by dialyses using a 10 kDa cutoff centrifugal filter unit from Millipore, Burlington, MA, USA), 150, 300, 450 and 600 mM. Samples were degassed by centrifugation (16.000xg per 2 minutes) before use. All experiments were done in duplicate.

For the detection of thermally induced amyloid formation presented in Figure 5, the 10 mg/mL protein solution used in the previous DSC experiments was used. The DSC-treated solution was first diluted two times in Buffer A and sonicated in ice for 30 seconds (5- and 25-seconds pulse on and off, respectively) using an amplitude of 10% in a BRANSON Digital Sonifire® (Soni-tech). Protein concentration was kept fixed at approximately 5 mg/mL in Buffer A.

### Size Exclusion Chromatography with Multi-Angle Light Scattering (SEC-MALS)

SEC-MALS measurements were performed on a miniDAWN TREOS multi-angle light scattering equipment with detectors at three angles (43.6°, 90°, and 136.4°) and a 659 nm laser beam (Wyatt Technology, CA). A Wyatt QELS dynamic light scattering module for determining hydrodynamic radius and an Optilab® rEX refractometer (Wyatt Technology) were used in line with a size exclusion chromatography analytical column (Superdex 200 HR10/300, GE Healthcare). BSA was used as a control sample. The protein solutions were eluted in a 50 mM Tris HCl, 500 mM NaCl buffer, pH 8.0, with a flow rate of 0.5 mL/min. The data were processed using ASTRA7 software (Wyatt Technology) with the following parameters: refractive index of 1.331, 0.890 cP for the viscosity of the solvent, and a dn/dc (refractive index increment) as 0.1850 mL/g (a common value for proteins). Protein solutions were centrifuged for 10 minutes at 10.000xg at a controlled temperature of 4°C before use.

### Circular Dichroism (CD)

Far-UV (195-260 nm) CD experiments were performed in a Jasco J-815 CD Spectrometer (JASCO Corporation, Easton, MD, USA) equipped with a Peltier temperature control and using a quartz cell with a 1-mm path length. The spectra were recorded with a scanning speed of 100 nm.min^-1^, spectral bandwidth of 1 nm, a response time of 0.5 s. The standard error presented were calculated from a triplicate sample measurement. All the protein stock solutions were at a minimum concentration where the dilution in a 20 mM sodium phosphate pH 8.0 + 0.03% DDM was at least 20-fold. All the experiments were performed at 25 °C.

### Bioinformatics

Protein intrinsic disorder predictions were performed using the PONDR VXLT, CAN_XT, and VS2L software (http://www.pondr.com/ accessed in 2020). The grand average of hydropathy (GRAVY) value for each modern and ancestor protein was calculated using ProtParam (https://web.expasy.org/protparam/ accessed in 2019) [^39^].

Coevolution and conservation analyses were performed using CONAN [^88^]. The analysis was performed over the Pfam [^80^] domain GRASP55_65 (PF04495), filtered by removing sequences not presenting at least 80% of the expected positions in its HMM and removing redundancy (80% identity). Coevolution signals were considered for residue-position pairs present in at least 5% of the sequences in the alignment with a minimum -log(p-value) of 10. The statistical validation method was based on Tumminello *et al*. [^89^], and the calculation was expanded to include marginally conserved residues.

Frustratrometer 2 (http://frustratometer.qb.fcen.uba.ar/ accessed in 2020) [^47^] was used to localize the level of frustration for each residue interaction [^47^]. The structural models of DGRASP55 (PDB ID 4KFW) and DGRASP65 (PDB ID 4KFV) were used in the analyses. For the ancestors, the molecular models of ANC1, 2, 3, and 4 were calculated by threading using AlphaFold2 [^90^], I-TASSER [^91^] and Swiss-model [^92^] (with GRASP55 - PDB IDs 4KFW as template).

The ConSurf Server was used to estimate the degree of evolutionary conservation in both mammalian DGRASP55 and DGRASP65 (PDB IDs 4KFW and 4KFV) [^48^]. The Homologues were collected from UniProt using HMMER, and Multiple Sequence Alignment was built using CLUSTALW. It was collected 300 homologues for each protein using a percentage of identity in the range of 30-95%.

## Acknowledgments

The authors thank the Fundação de Amparo à Pesquisa do Estado de São Paulo (FAPESP) (Grants No. 2015/50366-7 and 2012/20367-3), Conselho Nacional de Desenvolvimento Científico e Tecnológico (CNPq) (Grant no. 306682/2018-4), and Coordenação de Aperfeiçoamento de Pessoal de Nível Superior (CAPES) for the financial support. LFSM and MRBB thank FAPESP for the postdoctoral grants No. 2017/24669-8 and No. 2016/16328-3, respectively. MCM thanks FAPESP for the Ph.D. scholarship grant No. 2017/24669-8. EK thanks CAPES for the scholarship (88882.378790/2019-01). The authors also thank the Molecular Biophysics Group at Sao Carlos Physics Institute of the University of Sao Paulo for allowing access to the SEC-MALS (FAPESP Grant number: 15/16812-0).

## Supplementary Material

- Figures S1 to S9
- Table S1 and S2

