## Supplementary figures for "Resurrecting Golgi proteins to grasp Golgi ribbon formation and self-association under stress"

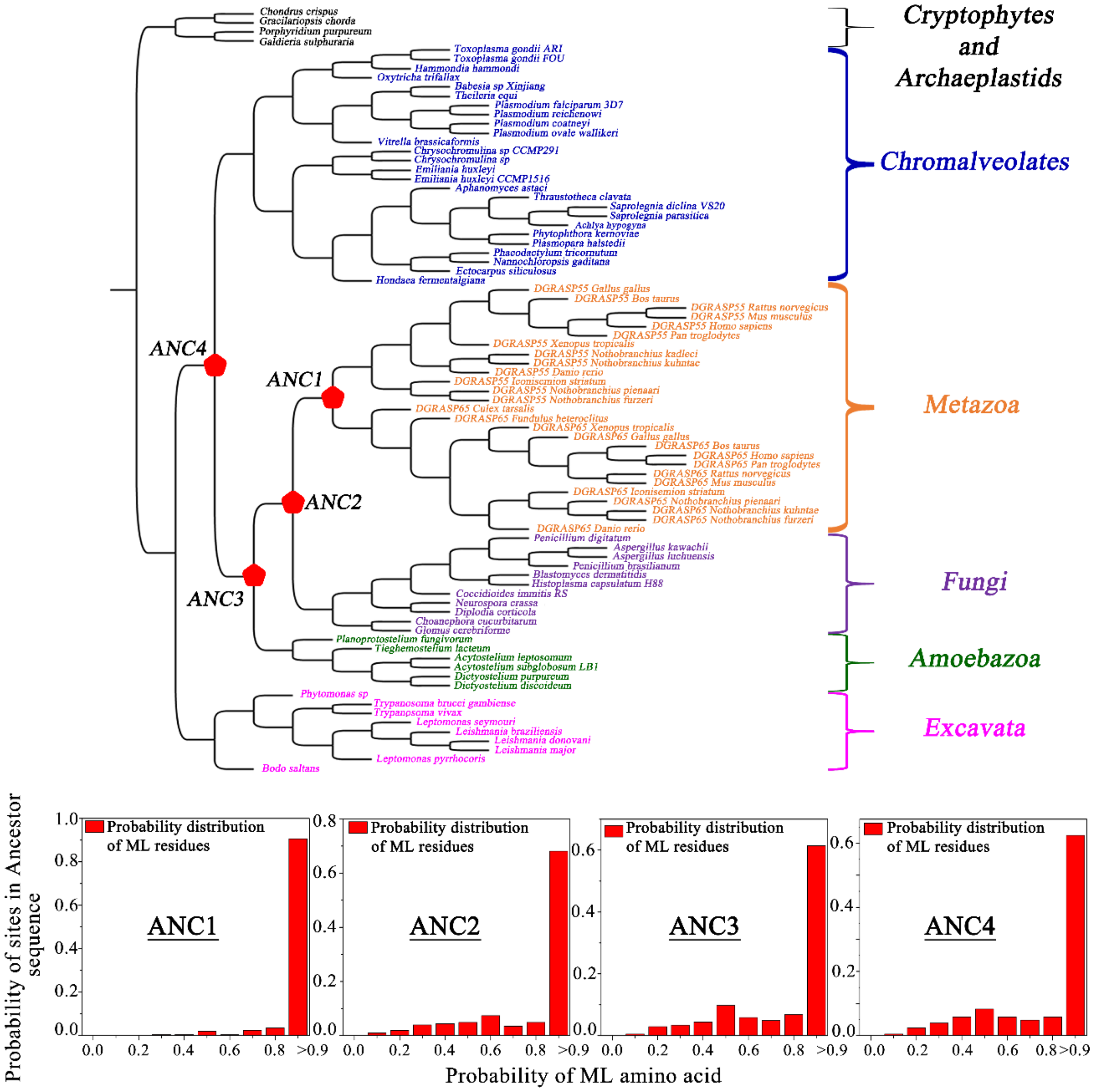


Figure S1: (A) Maximum likelihood phylogeny of eukaryotic GRASPs and the correspondent nodes where the ancestors were estimated. (B) Histogram of the posterior maximum likelihood probabilities.

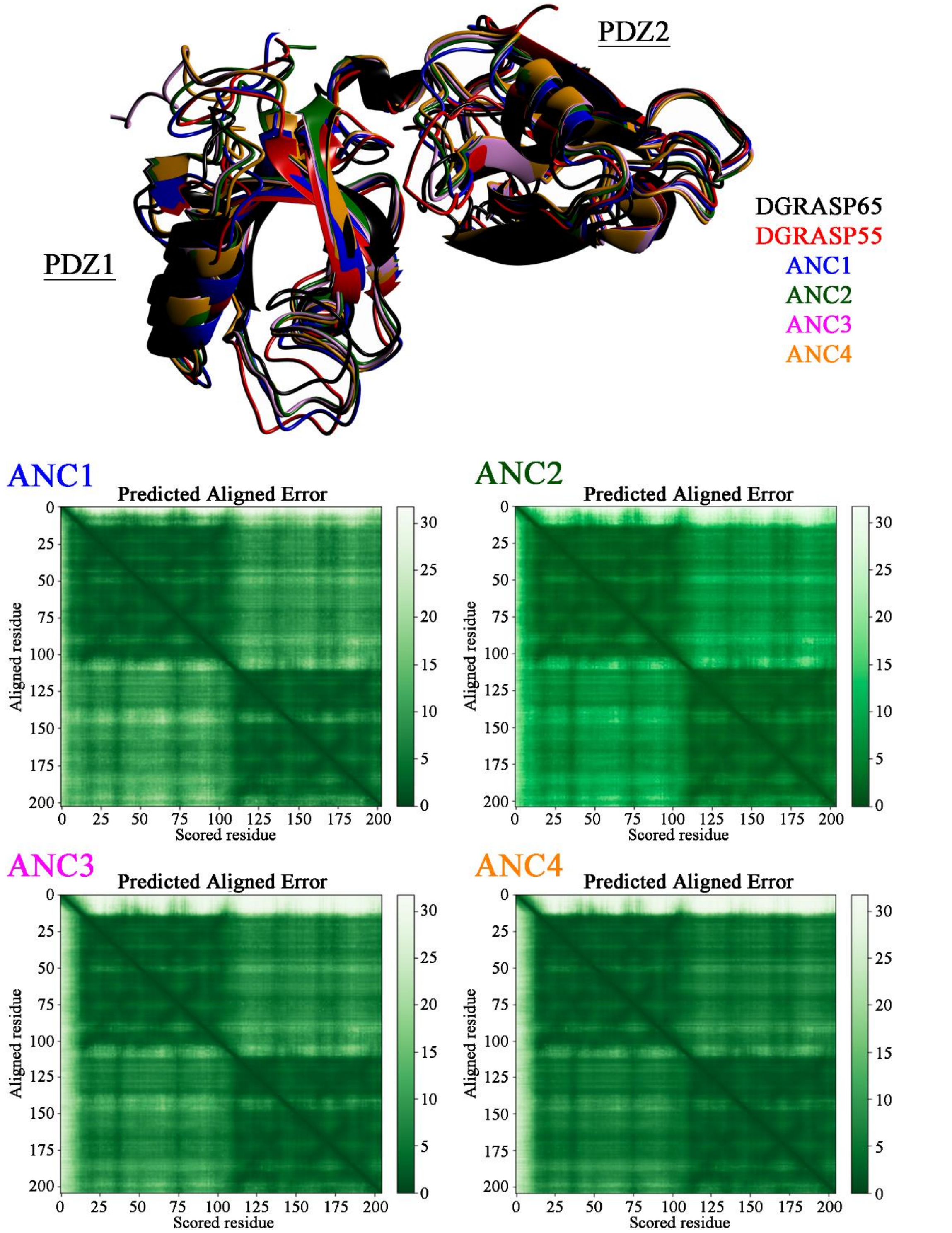


Figure S2: Structural superposition of the mammalian GRASPs (GRASP55 – PDB ID 4KFW and GRASP65 – PDB ID 4KFV) with their corresponding GRASP ancestors. Structural superposition was done using CCP4MG [1]. The ancestor GRASPs structural models were built using AlphaFold2 (DeepMind) [2] and the correspondent predicted aligned errors are shown. Except for the fully disordered N-terminal region, the major parts of the models have a per-residue confidence score higher than 90 (very high confidence).

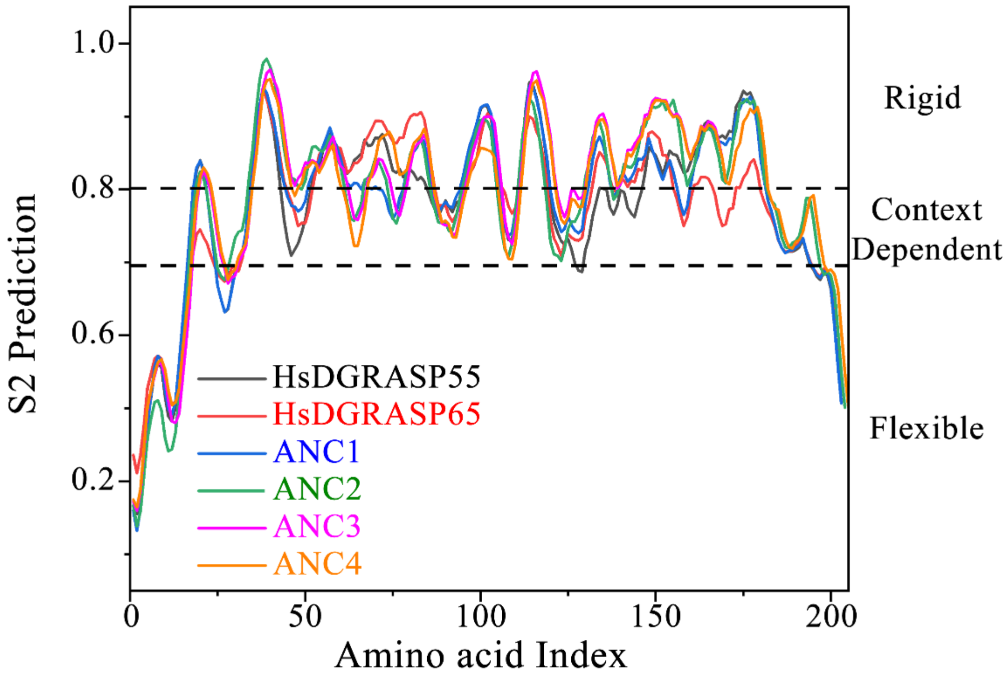


Figure S3: Prediction of protein backbone dynamics using DynaMine. Given a protein sequence, DynaMine predicts backbone flexibility at the residue level in the form of backbone N-H S2 order parameter values (http://dynamine.ibsquare.be/ accessed in 2021) [3,4].

(A) (B)

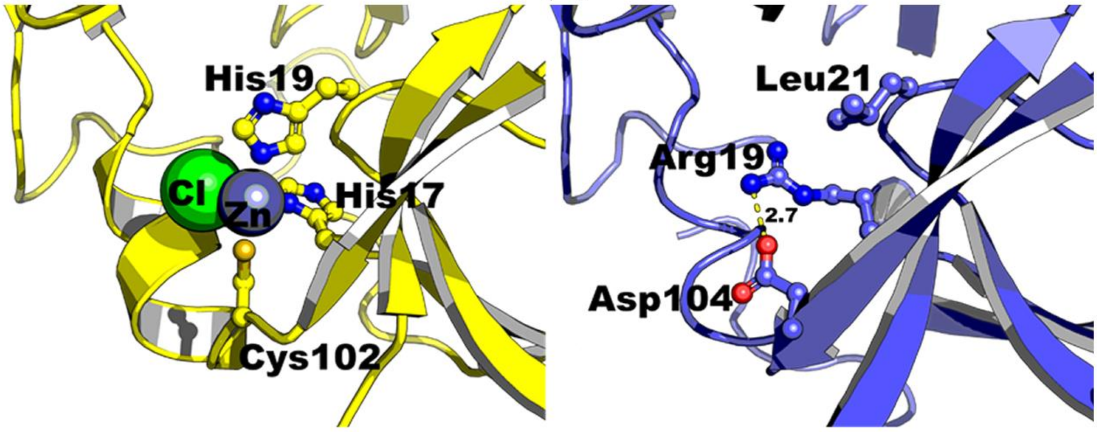


Figure S4: Putative structural changes in GRASP domains during evolution. (A) The zinc-binding site in GRASP65 (PDB ID: 4KFV). (B) A computational model of ANC4 with a salt bridge involving the structurally equivalent residues in the zinc-binding site.

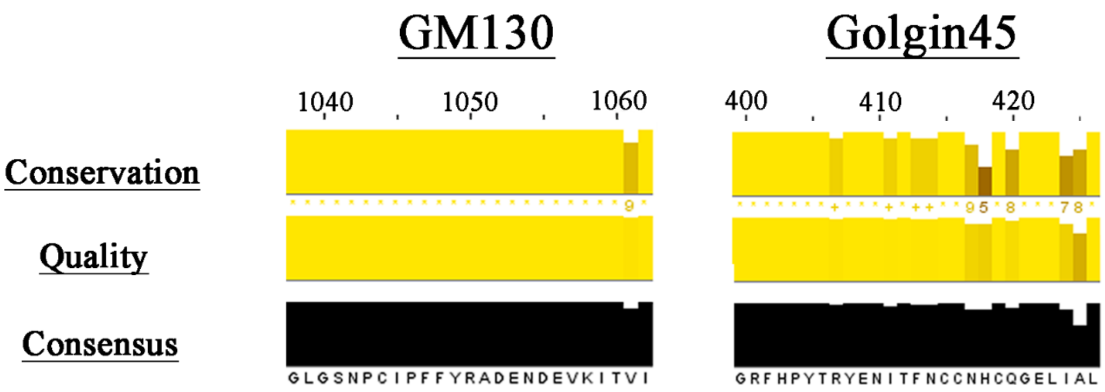


Figure S5. Analyses of the Multiple Sequence Alignment using Jalview [5]. 100 homologue sequences from the Non-Redundant Protein Sequence database of NCBI were collected for each Golgin and aligned using MUSCLE in MEGA-X. Only the GRASP binding region is presented. Protein sequences and the multiple sequence alignment can be provided upon request.

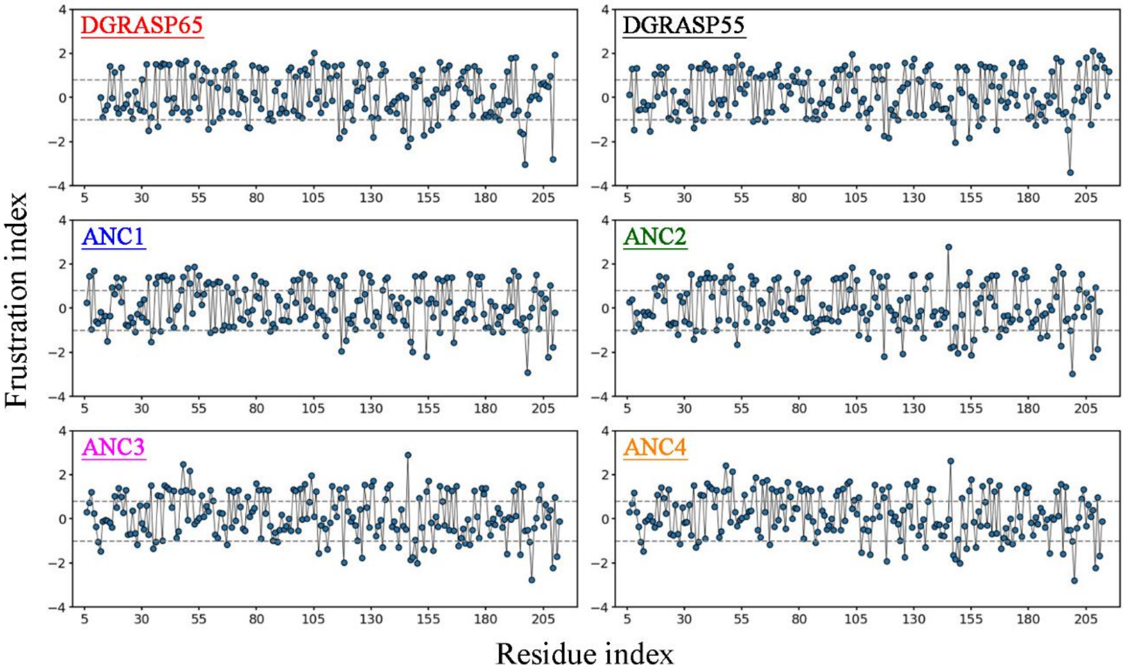


Figure S6. Degree of mutational frustration in mammalian GRASPs and their ancestors. The sites are considered highly or minimum frustrated if the local frustration index is lower than -1 or higher than 0.78, respectively.

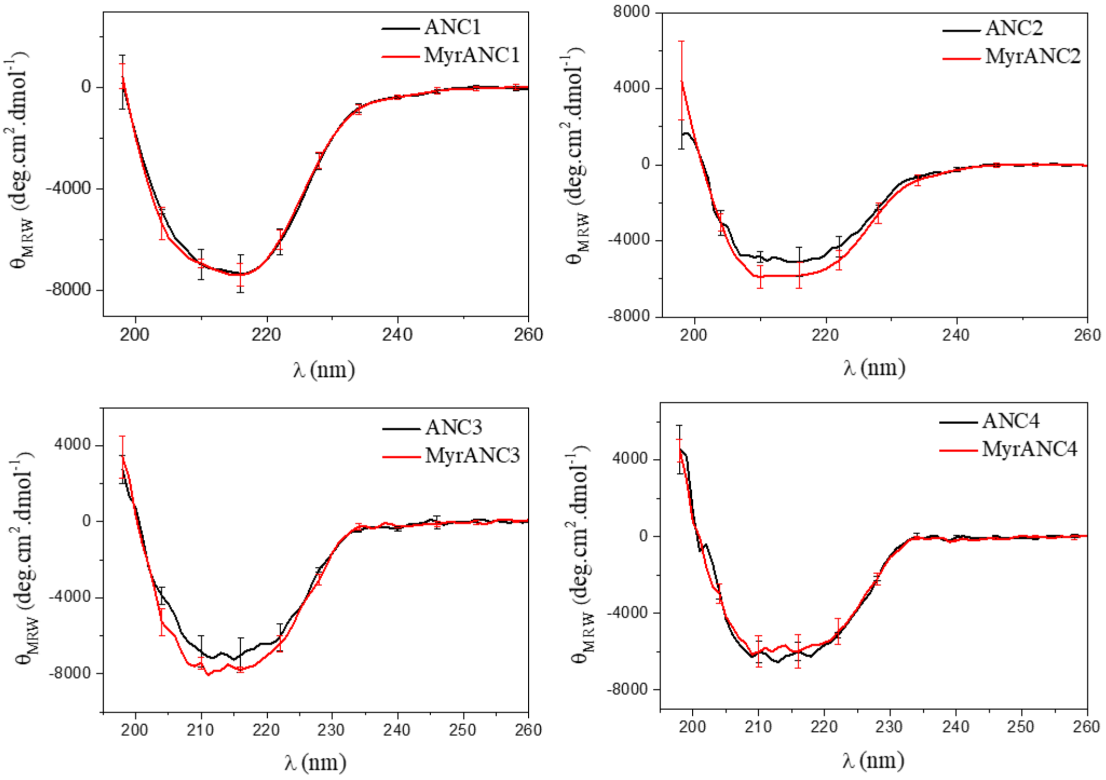


Figure S7: Circular Dichroism data of myristoylated and non-myristoylated GRASP ancestors in a detergent-containing buffer solution. The experiments were done in triplicate and at 25 °C.

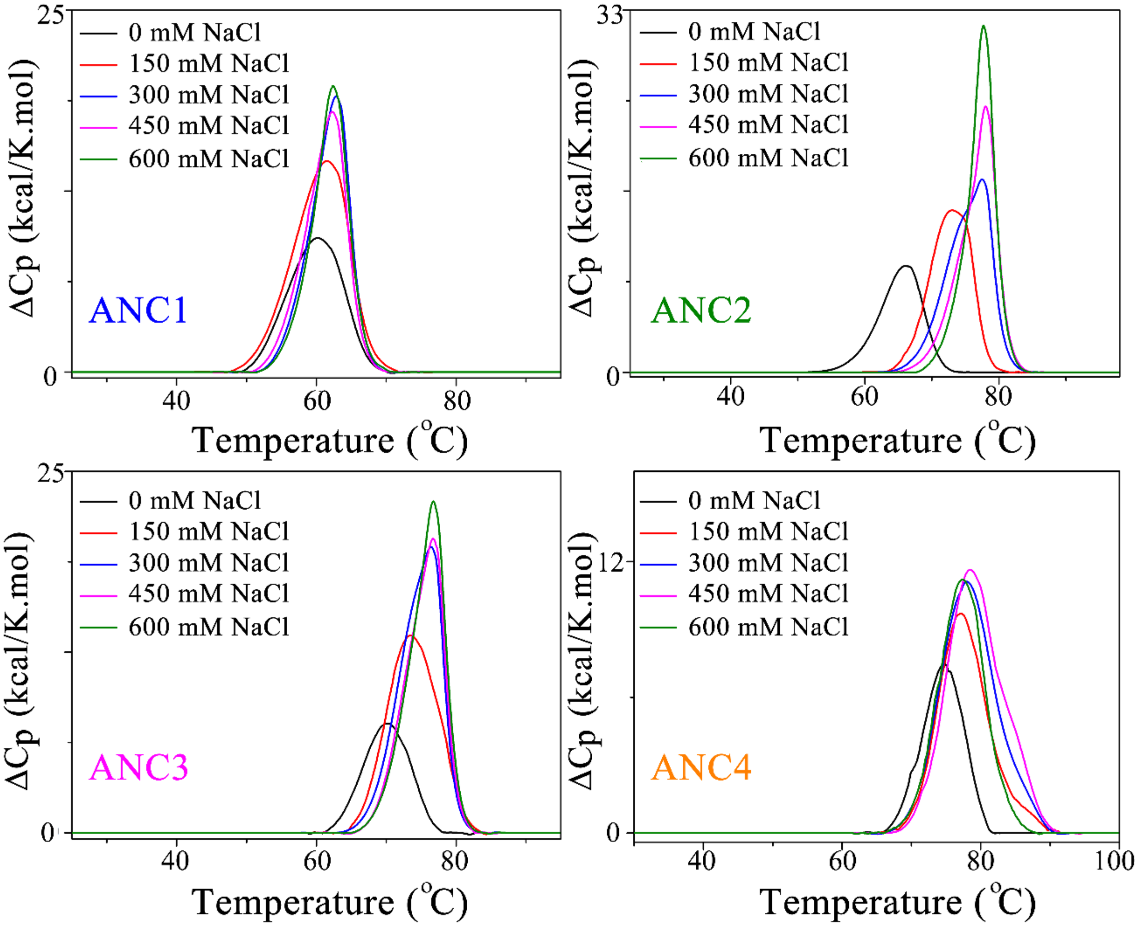


Figure S8. Excess heat capacity of the ancestor proteins at a fixed protein concentration of 2.5 mg/mL. The NaCl was varied from 0 to 150, 300, 450 and 600 mM in a 20 mM Tris/HCl pH 8.0 + 5 mM 2-Mercaptoethanol. The raw DSC traces were subtracted with the buffer baseline and then normalized by protein concentration.

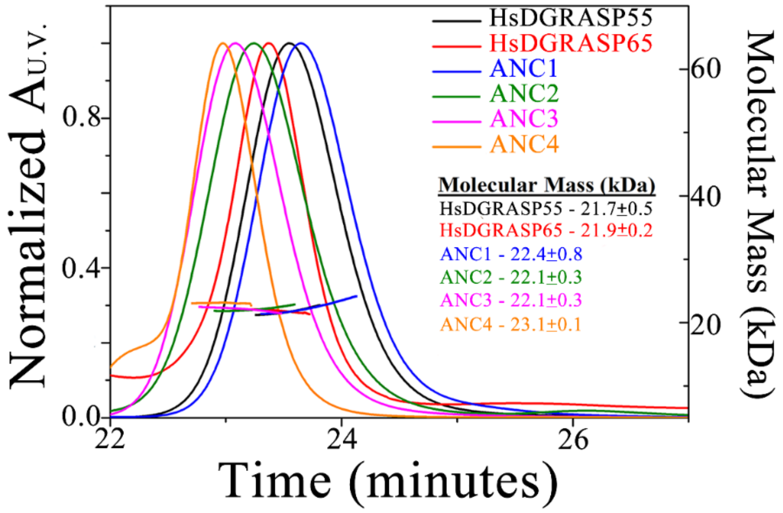


Figure S9: The HsDGRASPs and their ancestor's oligomerization were analysed by SEC-MALS in 50 mM Tris/HCl, pH 8.0, 500 mM NaCl. The calculated molecular mass of all the tested proteins, at this condition, correspond to a monomeric organization.

Table S1: N-terminal myristoylation of proteins by neural networks using Myristoylator [6] from Expasy:

| | Positive | Negative | Counters Positive/negative |
|---|---|---|---|
| *ANC1* | 0.876722 | 0.122739 | 23/2 |
| *ANC2* | 0.853763 | 0.145404 | 23/2 |
| *ANC3* | 0.992259 | 0.00789178 | 25/0 |
| *ANC4* | 0.992259 | 0.00789178 | 25/0 |

*ANC1/ANC2 are predicted to have medium confidence, while the prediction for ANC3/ANC4 have high confidence

Table S2: Thermodynamic parameters associated with the protein phase transitions.

| | ΔH (kcal/mol) | ΔS [kcal/(°C.mol)] | $T_M$ (°C) | $\Delta T_{1/2}$ (°C) |
|---|---|---|---|---|
| *ANC1* | 151.5 | 2.5 | 61.0 | 7.5 |
| *MyrANC1* | 138.5 | 2.2 | 62.0 | 8.5 |
| *ANC2* | 133.0 | 1.9 | 71.7 | 5.4 |
| *MyrANC2* | 87.4 | 1.3 | 68.4 | 5.4 |
| *ANC3* | 115.1 | 1.6 | 72.8 | 7.5 |
| *MyrANC3* | 95.8 | 1.3 | 74.5 | 7.6 |
| *ANC4* | 84.9 | 1.1 | 76.0 | 7.2 |
| *MyrANC4* | 102.1 | 1.4 | 75.0 | 7.7 |

The phase transition temperature, Tm, the calorimetric enthalpy change, ΔH, and the linewidth at half-height, $\Delta T_{1/2}$, were obtained from analysis of the thermograms with MicroCal Origin software. The entropy change, ΔS, at $T_M$ was calculated as ΔS = ΔH /Tm. Estimated uncertainties: ΔH (~5%), Tm (0.1°C), $\Delta T_{1/2}$ (0.1°C).
